# Structural Basis of Electron Transfer by the Human Nitric Oxide Synthase Holoenzyme Complex

**DOI:** 10.64898/2026.04.06.716795

**Authors:** Kanghyun Lee, Cristian Martinez-Ramos, Thomas H. Pospiech, Eric Tse, Miranda Lau, Yoichi Osawa, Daniel R. Southworth

**Affiliations:** Institute for Neurodegenerative Diseases, University of California, San Francisco, San Francisco, CA 94158, USA; Department of Biochemistry and Biophysics, University of California, San Francisco, San Francisco, CA 94158, USA; Department of Pharmacology, University of Michigan Medical School, 1301 MSRB III 1150 W. Medical Center Dr., Ann Arbor, MI 48109-5632, USA

## Abstract

Mammalian nitric oxide synthase (NOS) generates nitric oxide (NO), an essential signaling molecule in neurotransmission, inflammation, and cardiovascular regulation. NOS dysregulation contributes to neurodegeneration, septic shock, and ischemia. The structural basis of NOS function remains unclear, likely due to its dynamic, multi-component architecture. Catalysis requires electron transfer across five cofactors, with the oxygenase (Oxy) domain converting L-arginine to NO after receiving electrons from the reductase (Red) domain via calmodulin (CaM) activation. Here, we report cryo-electron microscopy structures of the human inducible NOS:CaM holoenzyme, revealing a previously unresolved architecture in which the Red:CaM arm spans the Oxy dimer in an apparent active state for cross-monomer electron transfer. The FMN subdomain is rotated ∼90° relative to the Red core, positioning its cofactor ∼9 Å from the heme, adjacent to a conserved aromatic residue within a narrow Oxy domain tunnel. Red–Oxy flexibility in the active state may facilitate electron shuttling among cofactors, defining how NOS cofactors are transiently aligned to enable catalysis.

## Introduction

Nitric oxide synthases (NOS) catalyze the formation of nitric oxide (NO) from L-arginine through a complex catalytic mechanism involving electron transfer among cofactors NADPH, FAD, FMN, and heme, with tetrahydrobiopterin (BH_4_) providing an additional electron required for oxygen activation during the two-step cycle^1,2^. In mammals, NO is a key signaling molecule that regulates diverse physiological processes, including neurotransmission, inflammation, and vasodilation^3,4^. Three conserved NOS isoforms mediate these functions: neuronal (nNOS), inducible (iNOS), and endothelial (eNOS), each exhibiting distinct expression patterns and modes of regulation^4^. Malfunction of NOS leads to aberrant NO production, which can react with superoxide to form highly reactive and toxic peroxynitrite^5,6^. Consequently, NOS dysregulation is implicated in many pathological conditions, including septic shock, cardiovascular conditions such as hypertension and stroke, and neurodegenerative disorders such as Alzheimer’s and Parkinson’s diseases^3,7,8^. Despite these critical roles in human health and decades of study, high-resolution structures of intact NOS enzymes have remained elusive, limiting understanding of the electron-transfer mechanism and hindering the development of therapeutics targeting disease-specific isoforms^9,10^.

NOS is active as a homodimer with each protomer composed of two core domains: an N-terminal oxygenase domain (Oxy), containing the dimer interface, heme and BH_4_, and a C-terminal reductase domain (Red), homologous to cytochrome P450 reductases, that consists of the NADPH/FAD (hereafter referred to as the FAD domain) and FMN subdomains (**Fig. 1a**)^1^. During catalysis, NADPH-derived electrons are transferred from FAD to FMN cofactors and then to ferric heme, enabling oxygen binding for the conversion of L-arginine to N-hydroxy-L-arginine and then to L-citrulline in the second step, releasing NO. NO generation is complex and proposed to require large domain rearrangements, including FMN subdomain dissociation from FAD, and FMN–Oxy domain interactions to enable electron transfer from the NADPH substrate to the stably bound redox cofactors FAD, FMN, heme, and BH_4_^2,11–13^. Heme insertion, regulated by molecular chaperones, is essential for dimerization^14–17^, which in turn is proposed to facilitate a trans mechanism of NO synthesis, wherein the FMN subdomain of one protomer transfers an electron to the heme moiety of the adjacent protomer^18–21^. However, structural evidence of inter-protomer interactions is currently lacking. Moreover, mammalian NOS isoforms are more complex, with NO synthesis activated by calmodulin (CaM) binding to a helical linker connecting the Oxy and Red domains^22–25^. This interaction induces conformational changes that transition the NOS reductase domain from a “shielded”, (referred to as Red^S^ or “input”) state for FAD-FMN electron transfer to a “de-shielded” (referred to as Red^DS^ or “output”) state, in which the FMN is proposed to rotate away from the NADPH-FAD center for electron transfer to heme in the adjacent Oxy domain^25–31^.

**Fig. 1.**
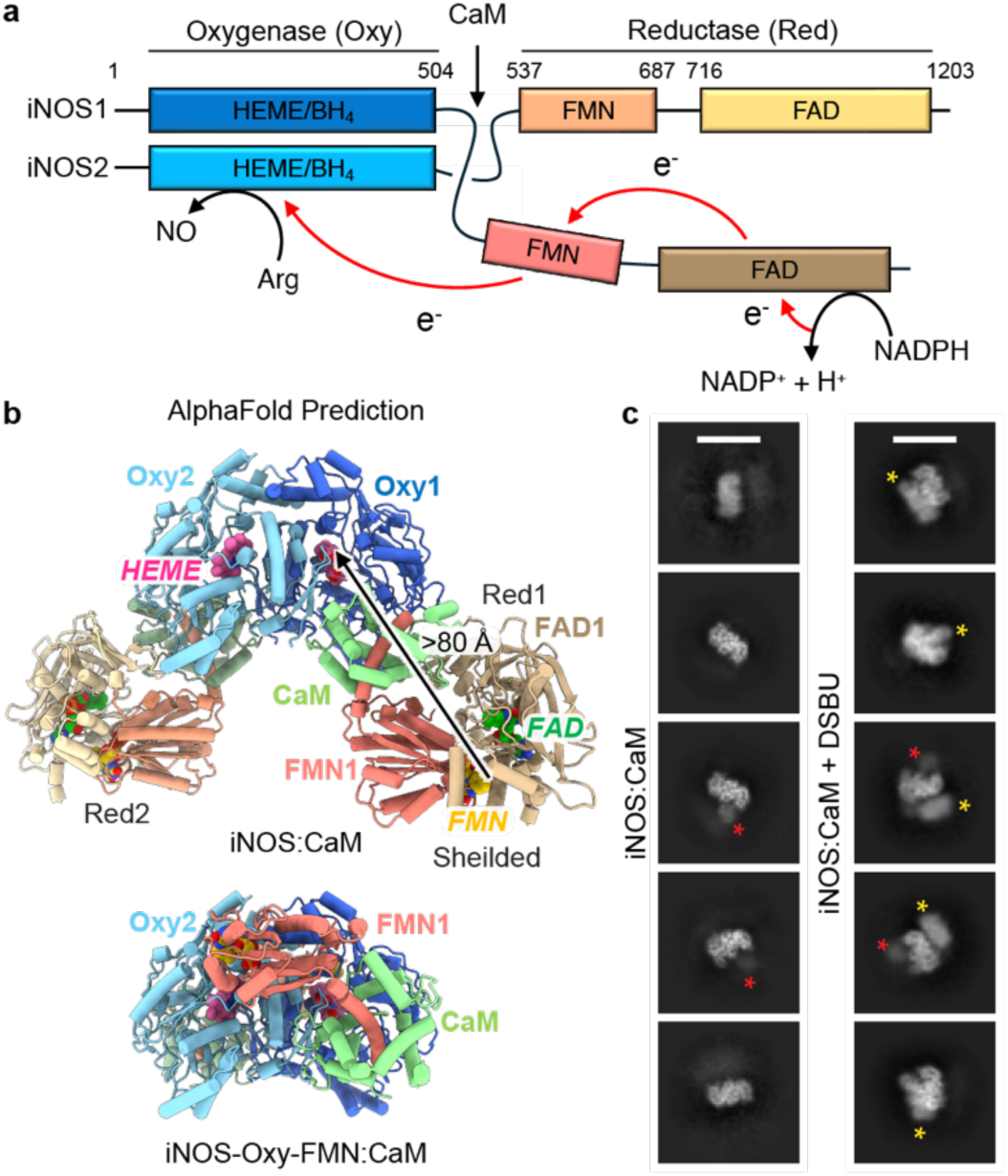
Domain architecture and 2D class averages of the iNOS:CaM complex. **a**, Schematic of the NOS homodimer, showing structured domains, CaM binding site, and electron transfer path. **b**, AlphaFold model of the full-length iNOS:CaM complex (top) and the complex without the FAD domain (bottom), predicted from human iNOS and CaM sequences (UniProt: P29475 and P0DP23, respectively). Flexible loops at the N-and C-termini are omitted for clarity. **c**, Representative 2D cryo-EM class averages of the iNOS:CaM complex, without (left) or with (right) DSBU, with poorly resolved (red asterisk) and well-resolved (yellow asterisk) Red-CaM density indicated (scale bar = 120Å).

Currently, high-resolution structural information is limited to NOS domain truncations. Structures of the NOS Oxy domain identify that the catalytic site adopts a distinct α/β “catcher’s mitt” fold that houses the heme and BH₄ binding sites adjacent to a hydrophobic dimer interface, illustrating how binding of these cofactors promotes assembly of the holoenzyme^32–34^. The highly conserved substrate-binding site is located adjacent to heme and has been targeted for the development of NOS inhibitors, with multiple inhibitor-bound structures available for the Oxy dimer^35^. The Red domain structure determined for nNOS identifies buried cofactor sites in an FAD-FMN shielded conformation (Red^S^)^36^. In contrast, comparison to the structure of the iNOS FMN subdomain bound to CaM indicates how the FMN could rotate away from the FAD upon CaM binding^25^. We previously identified CaM-dependent opening of the Red domain in intact nNOS complexes, suggesting a shielded-to-de-shielded switch^37^. While these and other low-resolution EM studies established that large CaM-dependent changes in the FMN position can occur, the NOS:CaM complex is conformationally dynamic, and Red-Oxy interactions may only be transiently sampled during catalysis^37–39^. More recently, crosslinking mass spectrometry (CXL-MS) coupled with AlphaFold Multimer prediction studies has captured FMN-Oxy interdomain contacts, leading to compelling structural models for how the FMN domain may be positioned within 15 Å of the heme pocket^40–42^. These FMN-Oxy interaction models are consistent with previous studies employing complementary mutagenesis^30,31^, electron paramagnetic resonance spectroscopy^43–45^, hydrogen-deuterium exchange^26^, and molecular dynamics simulations^28^, supporting the trans-electron-transfer mechanism that occurs within a feasible distance^20,21^. Nonetheless, the structural basis of the de-shielded state and interdomain electron transfer remains undefined.

These fundamental gaps in understanding prompted us to focus on the structural characterization of human iNOS, which binds constitutively to CaM and lacks additional flexible regulatory or autoinhibitory elements found in other isoforms^46–49^. Moreover, we sought to apply our previous chemical crosslinking techniques to trap Red-Oxy interactions for cryo-EM^40^. With these efforts, we determined structures of the iNOS:CaM complex to an overall ∼3.0 Å resolution that capture the interdomain electron transfer state, revealing the Red domain adopts an FMN-rotated, de-shielded conformation that contacts the Oxy domain, positioning the exposed FMN cofactor directly over the heme binding site of the adjacent protomer. This architecture is maintained by distinct intra-and interprotomer and CaM interaction sites that we identify as required for activity. 3D classification analysis reveals Red-Oxy conformational changes that enable cofactor alignment and the formation of an interdomain electron-transfer tunnel, coordinated by a conserved hydrophobic network, which orients the FMN within 9 Å of the active-site heme. Based on two distinct positions of the Red domain, we propose that flexible contact with the Oxy dimer enables electron transfer to favor FAD-FMN in one state and FMN-heme in a second state, revealing a conformationally guided reaction cycle for NO synthesis. Together, this work provides the long-sought structural basis for the interdomain electron transfer state of NOS enzymes, offering fundamental insights into the catalytic mechanisms conserved across the cytochrome P450 superfamily and revealing new interaction interfaces for potential isoform-specific therapeutic development.

## Main

### Crosslinking captures iNOS:CaM Red-Oxy interaction state for cryo-EM

Here, we focused on characterizing the structure of human iNOS, based on the hypothesis that this isoform may more readily adopt the Red-Oxy electron transfer state compared to the larger nNOS and eNOS isoforms^50^. Notably, the predicted structure, using AlphaFold3, shows the full-length iNOS:CaM tetramer in an inactive extended conformation with the FMN and heme separated by more than 80 Å and the Red domain in a shielded state, similar to that determined crystallographically, but with a slightly altered orientation (**Fig. 1b, top**)^36^. Intriguingly, the predicted arrangement of a truncated iNOS–CaM complex lacking the FAD domain shows the FMN domain docked on the surface of the adjacent Oxy monomer in a position that appears competent for electron transfer (**Fig. 1b, bottom**). This is mediated by CaM, which engages the Oxy domain and binds the CaM-binding helix to position FMN, as in the previous FMN-CaM crystal structure^25^. Cross-linking mass spectrometry (XL–MS) studies support this prediction, yet the intact arrangement for electron transfer, including FAD-FMN positioning remains unclear ^40–42^. These observations underscore the limitations of structure predictions based on domain truncations and in the context of full-length complexes.

To characterize the iNOS:CaM complex, proteins were co-expressed in *E. coli* and purified using a tandem affinity strategy with Ni-NTA and 2’,5’-ADP resin (**Extended Data Fig. 1a**)^51,52^. The iNOS:CaM tetramer complex, a dimer of iNOS:CaM dimers (hereafter referred to as the dimer), was isolated by size-exclusion chromatography (SEC) and subsequently verified to have the expected molecular weight of ∼290 kDa for the dimer by multi-angle light scattering and XL analysis (**Extended Data Fig. 1b–d**). Heme incorporation (∼2.6 nmol P450 mg⁻¹ NOS) and NO synthesis activity (iNOS activity ≈ 94 nmol NO min⁻¹ nmol⁻¹ P450) were confirmed by CO binding and oxyhemoglobin assays, respectively, establishing formation of an active holoenzyme complex (**Extended Data Fig. 1e, f**)^53^.

Our previous cryo-EM analysis of nNOS:CaM (with and without XL) yielded 2D averages that exhibited Oxy dimer density but no additional density for CaM or the Red domains^40^. Initial 2D classification analysis of iNOS:CaM similarly shows well-resolved density for the Oxy dimer with only weak additional density in certain averages, indicating CaM and the Red domain remain flexible for iNOS (**Fig. 1c (left)**). We therefore incubated iNOS:CaM with disuccinimidyl dibutyric acid (DSBU) prior to sample vitrification for cryo-EM to increase stability and potentially capture Red-Oxy interactions (See Methods). Strikingly, 2D averages of XL iNOS:CaM show additional globular density adjacent the Oxy dimer indicative of the Red:CaM bound to the Oxy dimer (**Fig. 1c (right), yellow asterisk**). Density is limited to one face of the dimer, with only weak CaM density on the opposite side (**Fig. 1c (right), red asterisk**), suggesting that the second Red:CaM arm remains flexibly linked and therefore cannot be resolved. These results reveal that XL procedures stabilize the Red:CaM relative to the Oxy dimer in an arrangement not previously observed in related structural studies^37–40^. Moreover, the iNOS high-affinity interactions with CaM and absence of regulatory elements may provide further stability for cryo-EM structural analysis with XL compared to other isoforms.

### The iNOS:CaM structure reveals Red-Oxy interactions in an FMN-heme electron transfer state

Here, we aimed to determine high-resolution structures of the iNOS:CaM complex. To overcome preferred orientation and conformational variability we employed detergent addition, tilt collection, and extensive 3D classification strategies to generate classes containing clear Red-CaM density (**Extended Data Fig. 2**). Heterogeneous refinement yielded two major resolvable classes: a predominant class (containing ∼35% of the data) exhibiting a well-defined Oxy dimer with only weak CaM density, while a second dominant class (containing ∼21% of the data) shows strong density for one Red:CaM arm bound to the Oxy dimer (**Extended Data Fig. 2b–d**). Refinement of the Oxy dimer class resulted in a 2.6 Å resolution map and, following atomic modeling, we identified the structure closely matches previous crystal structure (ɑ-carbon RMSD between 409 pruned atom pairs = 0.5 Å compared to murine iNOS (PDB: 3NQS)) and includes well-resolved density for heme and BH_4_ cofactors (**Extended Data Fig. 3d and 4a–d**). Visualization of the unsharpened, filtered map exhibited no additional density for the Red beyond the flexibly associated CaM indicating these domains are likely not engaged with the Oxy dimer in this class (**Extended Data Fig. 4e**). The class containing Red:CaM density refined to an overall 3.0 Å resolution for the consensus map and local resolution estimates indicate high, ∼2.5 Å resolution for the Oxy dimer while the Red:CaM arm is at a more modest ∼4.5–6.5 Å resolution, presumably due to its flexibility relative to the Oxy dimer (**Fig. 2a, Supplementary Video 1, and Extended Data Fig. 3a**). A complete atomic model was built by separately docking the AlphaFold-predicted Oxy dimer, an FAD subdomain, and the CaM-bound FMN subdomain structure (PDB: 3HR4) followed by refinement using RosettaRelax^54^ and ISOLDE^55^ (**Fig. 2b**).

**Fig. 2.**
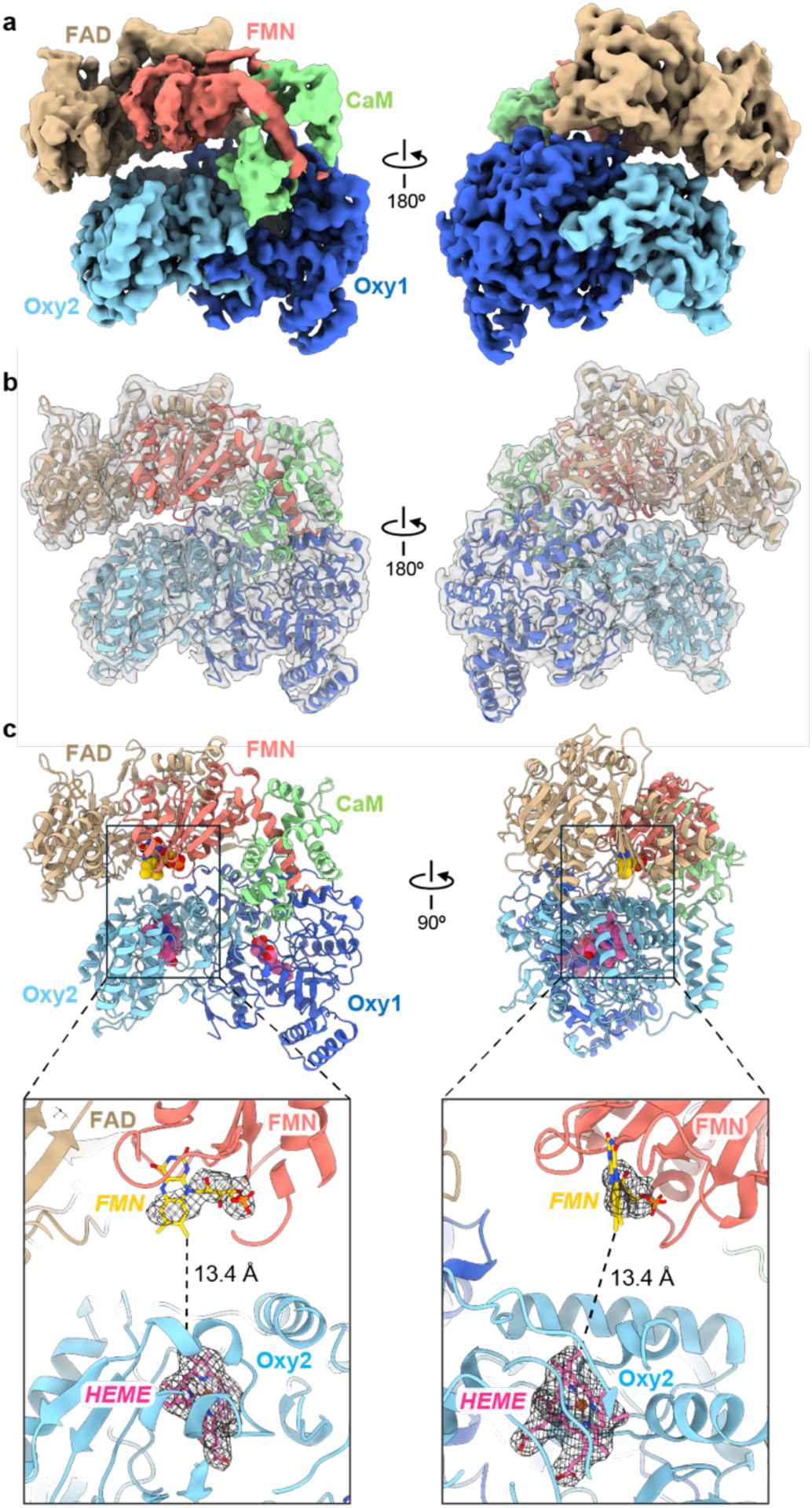
Cryo-EM map and model of iNOS:CaM in an electron-transfer-competent engaged state. **a**, Final consensus map of the iNOS:CaM complex at 3.0 Å resolution, segmented and colored by domain. **b**, Map and refined model of iNOS:CaM, colored by domain with cryo-EM density in transparent grey. **c**, Model of iNOS:CaM, with FMN (yellow) and heme (pink) shown in space fill and enlarged views (lower panels) of the FMN-heme arrangement, showing the cofactor distance and difference map density in mesh (calculated as the cryo-EM map – the cofactor-free model).

**Fig. 3.**
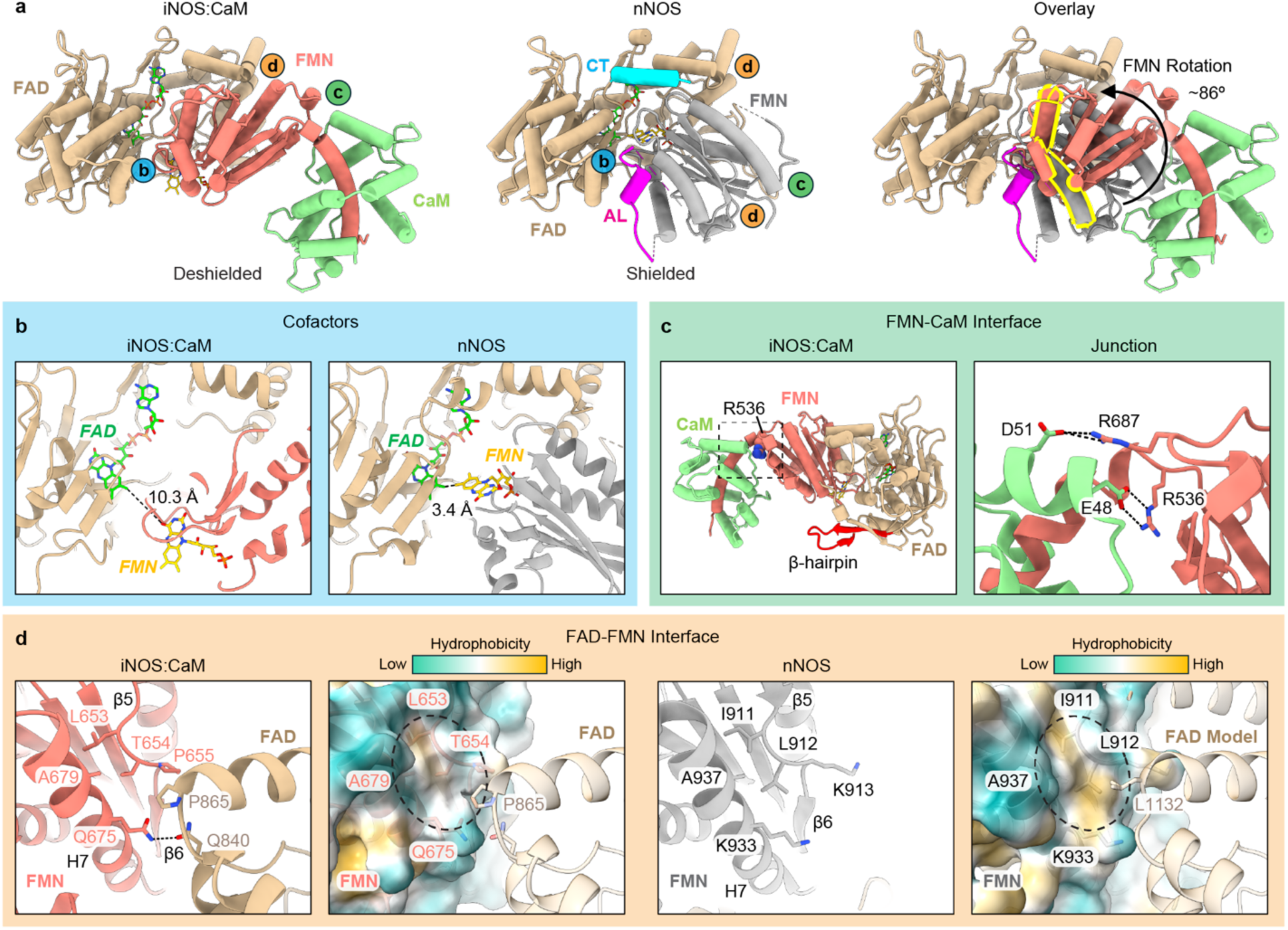
Reductase domain and FMN adopts a deshielded, output state in the iNOS:CaM structure. **a**, The Red domain conformation and FMN subdomain rotation to the deshielded state are shown by comparing Red:CaM from the iNOS structure (left, determined here) to the nNOS truncated Red domain structure (middle; PDB: 1TLL), with the FMN (grey) in a shielded conformation^36–39^, and the nNOS-specific CT (teal) and AL (magenta) motifs shown. An overlay (right) of the iNOS Red:CaM (colored) and nNOS Red (FMN in grey), aligned to the respective FAD subdomains, shows an ∼86° rotation of the FMN, with helix α5 highlighted (yellow outline) to illustrate the change. AL: autoinhibitory loop; CT, C-terminal tail. **b**, Enlarged view of the FAD-FMN subdomain interface for iNOS:CaM (left) and nNOS (right), showing a large (10.3 Å) FAD-FMN cofactor distance in iNOS:CaM, consistent with the deshielded, compared to the smaller (3.4 Å) distance observed in nNOS. **c**, The iNOS FMN-CaM interaction interface is shown (left) with an enlarged view (right) of the CaM-stabilized FMN hinge and key interaction residues. **d**, The de-shielded-specific FAD-FMN interface in iNOS:CaM with a hydrophobicity surface map shown with key interacting residues (left panels), in comparison to nNOS and a hydrophobicity map of nNOS in a hypothetical de-shielded state and potential conserved interactions (right panels).

**Fig. 4.**
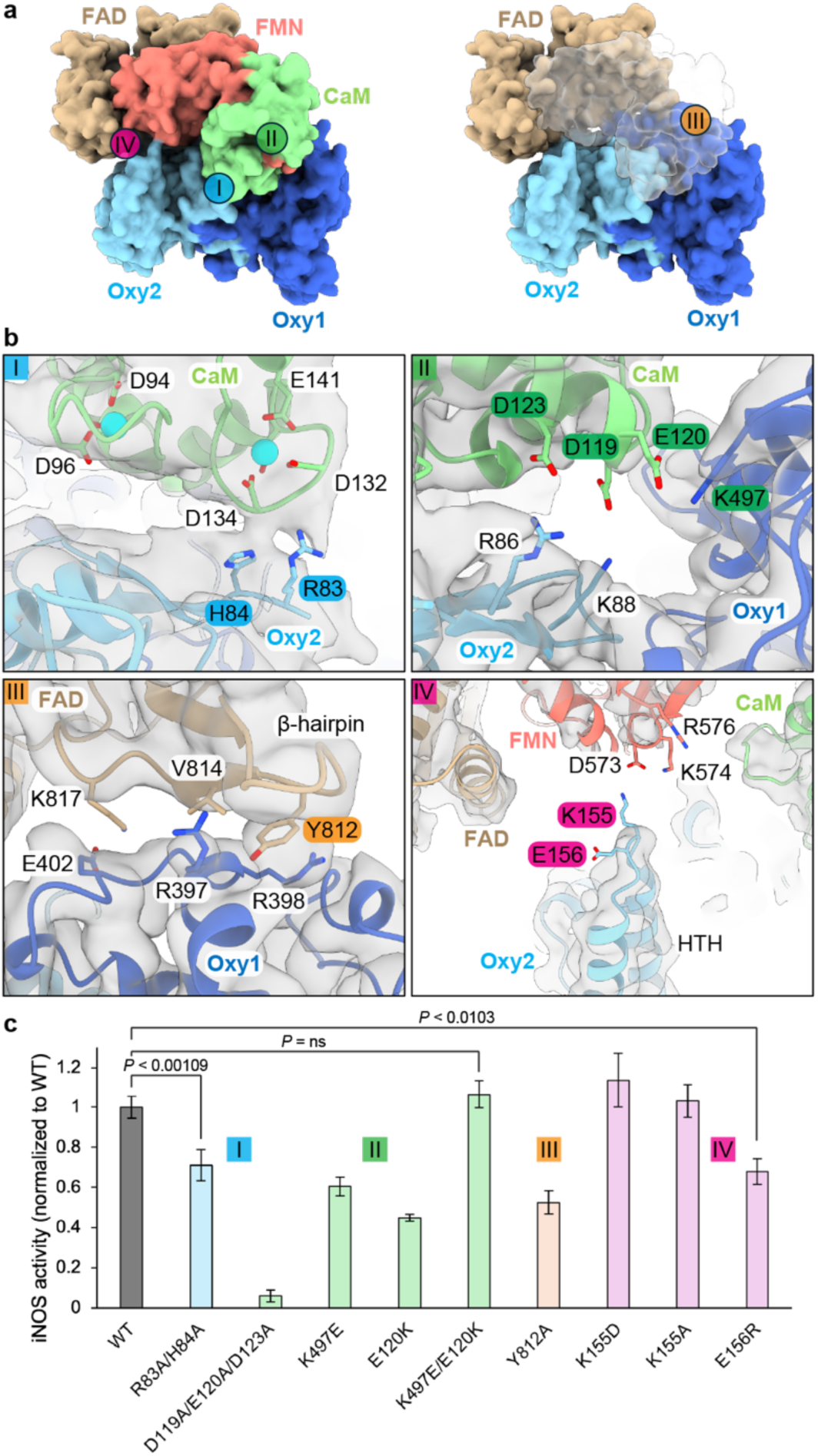
Intermolecular iNOS:CaM contacts stabilize the engaged state for NO synthesis. **a**, Surface model structure of iNOS:CaM, with four distinct interaction interfaces indicated (I-IV) for further examination (below). **b**, Enlarged map–model views of interprotomer and interdomain contacts: (I) Oxy2:CaM, (II) CaM:Oxy dimer, (III) FAD:Oxy1, (IV) and FMN:Oxy1 (IV) with interacting residues targeted for mutagenesis indicated by color. **c**, Mutagenesis and NO generation activity analysis of interaction site residues, colored by site as in (b). NOS Activity (Y-axis) measured by the conversion of oxyhemoglobin to methemoglobin and standardized to the P450 concentration, with the mutant activity shown normalized to WT level. Data represent mean ± SD of three technical replicates. *P*-values were determined by a one-sided paired two-sample t-test, assuming normality of differences, with no adjustment for multiple comparisons.

From the atomic model, we find that iNOS:CaM adopts an “engaged” conformation in which the Red domain extends across the Oxy dimer, positioning a deshielded FMN subdomain next to the heme in the Oxy domain of the second protomer (referred to as Oxy2) (**Fig. 2c, top**). Based on difference map analysis we identify strong density for the FMN and heme cofactors, revealing the FMN is exposed, projecting towards the Oxy surface, and within 14 Å of the heme, a distance proposed to be sufficient for electron transfer across the domains (**Fig. 2c, bottom**)^13,27^. We note the density corresponding to the FMN cofactor is weaker than that of the heme, consistent with the flexibility of the Red domain and potential conformational variability of the FMN subdomain (**See Below and Fig. 5 and Extended Data Fig. 7**). Overall, this structure captures the active FMN-to-heme electron-transfer state required for NO synthesis, demonstrating that NO production occurs *in trans*, across the monomers and depends on the dimeric architecture, consistent with prior biochemical evidence^20,21^.

**Fig. 5.**
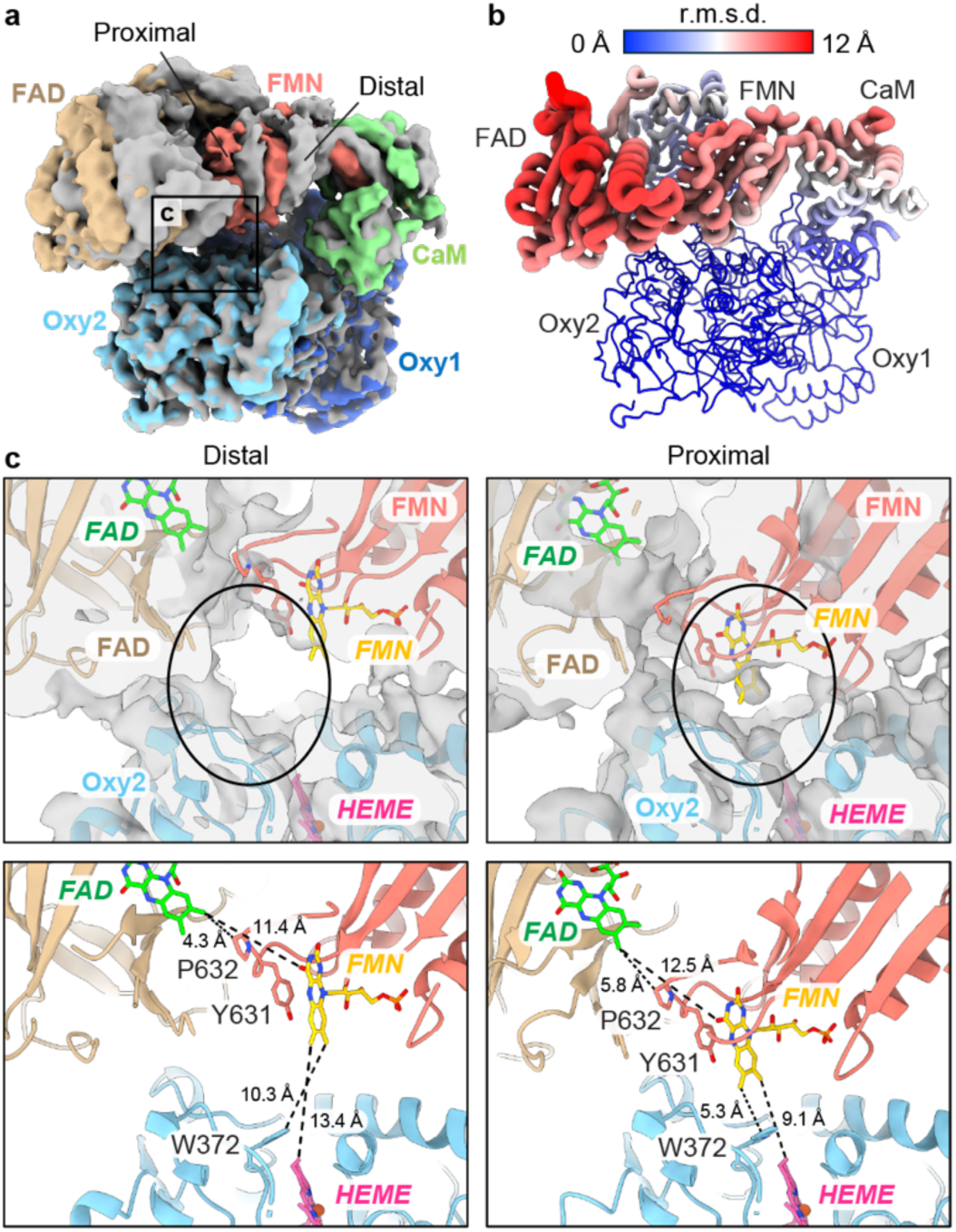
Conformational flexibility of the Red:CaM arm facilitates cofactor alignment for inter-domain electron transfer. **a**, Overlay of distal (gray) and proximal (colored) iNOS:CaM refined maps, identified from 3DVA and aligned to the Oxy dimer, showing relative shifts in the Red:CaM position. **b**, The iNOS:CaM proximal state colored according to Cα r.m.s.d. values relative to the distal state (aligned to the Oxy dimer). The largest changes (>10 Å, shown in red with wider tubes) are identified for the FAD and FMN subdomains. **c**, Enlarged views of the FAD–FMN–heme cofactor site in distal (left) and proximal (right) conformations. Map+model (top) and model (bottom) views show convergence of the FMN–heme position in the proximal state relative to the distal state (circle). Distances between cofactors and conserved electron-transfer residues are indicated (bottom).

### The Red:CaM adopts an FMN de-shielded output state to engage the Oxy dimer for electron transfer from FMN to heme

The structure suggests that two separate events occur: (1) FMN deshields from FAD, and (2) the deshielded Red domain with CaM interacts with the Oxy dimer to form the engaged state required for interdomain electron transfer. To define the de-shielded state, we first sought to characterize the conformation and interactions of the Red:CaM arm (**Fig. 3**)^36^. We compared the Red:CaM arrangement in this structure with the nNOS Red-domain crystal structure, which serves as a benchmark for the shielded arrangement (Red^S^), in which the FMN subdomain is docked against the FAD subdomain (**Fig. 3a, middle**). In this conformation, FAD–FMN electron transfer is supported, but transfer to heme in the Oxy domain is considered unlikely due to the buried arrangement of the FMN cofactor^36^. By aligning the FAD domains, we identify that the FMN is rotated nearly 90° in the iNOS:CaM structure, which results in a substantial repositioning of the FMN cofactor towards the Oxy domain (**Fig. 3a, right, and Supplementary Video 2**). This rotation yields a distinct cofactor arrangement and new interaction interfaces that we propose define the de-shielded Red conformation for iNOS (Red^DS^), which is required for the Oxy dimer interaction to form the engaged state and enable electron transfer from FMN to heme.

Notably, the Red^S^ conformation is not compatible with the iNOS:CaM map, as evidenced by clashes between the FMN and Oxy domains (**Extended Data Fig. 5a**). Conversely, when we separately fit the nNOS FAD and FMN subdomains into our iNOS Red^DS^ map we identify only minor clashes would occur for the autoinhibitory loop (AL) and C-terminal tail (CT) (absent in iNOS) with the deshielded FMN orientation, indicating that this FMN rotation could be accommodated in other isoforms with slight rearrangement of these flexible regulatory elements (**Extended Data Fig. 5a**). Thus, we conclude that the Red^DS^ conformation we observe for iNOS maybe conserved among the three isoforms and required for Red engagement of the Oxy domain. In the Red^DS^ conformation both the FMN and FAD cofactors become solvent exposed, separated by ∼10.3 Å compared to ∼3.4 Å in the Red^S^ state (**Fig. 3b and Extended Data Fig. 5b**). Although this longer distance may be compatible with FAD–FMN electron transfer^12,13,56^ the FMN isoalloxazine ring is rotated ∼90°, such that the C7 and C8 methyl groups are no longer juxtaposed with those of the FAD and instead project toward the Oxy domain (**Fig. 2c and 3b**). This FMN orientation is also observed in the structure of the bacterial FMN-heme cytochrome P450BM-3^57^ and thus indicates a state that supports interdomain electron transfer to the Oxy domain.

### CaM, FMN, and FAD interaction network supports the deshielded Red state

Upon inspection of the CaM interaction, we identify that the FMN rotation is supported through the previously characterized “hinge region”^25^ that includes salt bridge contacts between CaM E48 and iNOS R536, which reside at the junction between the CaM binding helix and the FMN subdomain **(Fig. 3c and Extended Data Fig. 5c)**. This salt bridge interaction matches that observed in the FMN:CaM crystal structure^25^, and was previously determined to be necessary for CaM-activated NO production^58^. Additionally, we identify a putative salt bridge between D51 of CaM and R687 in the FMN domain, which appears to further support the hinge and overall Red^DS^ conformation. Intriguingly, previous analysis of the FMN:CaM structure identified four slightly different positions of the CaM-binding helix relative to the FMN domain, based on flexibility at the R536 hinge site^25^. In the context of the intact iNOS:CaM complex, we observe the CaM-binding helix to be further shifted by an additional ∼6° rotation relative to the FMN domain (**Extended Data Fig. 5d (left))**. Importantly, when the CaM-binding helix is instead aligned between these structures to compare positioning relative to the FAD subdomain we identify the FMN cofactor is shifted by as far as ∼15 Å from the previous FMN:CaM structure, positioning the FMN cofactor closer to the Oxy domain compared to what can be modeled from the FMN:CaM structure (**Extended Data Fig. 5d, middle and right panels**). Thus, in the intact iNOS:CaM complex, CaM binding causes a larger FMN shift toward the Oxy domain than previously anticipated, through additional interdomain contacts, further establishing its requirement for reorienting the FMN for NOS activity. We note the CaM-FMN orientation in our structure is incompatible with the FMN in the shielded conformation, primarily due to a steric clash that would occur between the β-hairpin finger motif formed by β11 and β12 of the FAD domain (following the previous numbering system for the Red^20^) and the C-lobe of CaM (**Extended Data Fig. 5e**)^25^. Thus, considering that CaM is thought to be constitutively bound to iNOS, the canonical Red^S^ state may not be readily sampled for this isoform.

Finally, we identify unique contacts between the FAD and FMN subdomains that were not previously observed and are likely to help stabilize the Red^DS^ state (**Fig 3d**). Specifically, a helical turn between β5 and β6, together with an adjacent helix H7 of the FMN domain, forms a hydrophobic pocket that contacts residue P865 in the FAD domain through potential hydrophobic interactions with T654 and P655. In contrast, the adjacent Q840 residue in the FAD appears capable of forming a hydrogen bond with Q675, further stabilizing the interaction (**Fig. 3d, left panels**). We note that the surface architecture and hydrophobicity are conserved in human NOS isoforms. Thus, we predict that this interaction may occur between nNOS and eNOS to stabilize their respective Red^DS^ states (**Fig. 3d, right panels**). Overall, characterization of the Red:CaM portion of the iNOS structure reveals distinct FMN:CaM and FAD:FMN interactions that define and stabilize a Red^DS^ state, which is required to engage the Oxy dimer and is likely conserved across isoforms.

### Distinct interdomain interactions define the engaged state of the iNOS:CaM complex

We next aimed to define Red:CaM interactions with the Oxy dimer that enable FMN engagement with the adjacent protomer (Oxy2). Overall, we observe 4 distinct interdomain and interprotomer interaction sites that constitute the Red-Oxy engaged arrangement (**Fig. 4a**). Upon inspection of CaM interactions, we identify substantial charge complementarity between the highly acidic surface of CaM and two distinct basic regions on the Oxy dimer (**Fig. 4b, I–II and Extended Data Fig. 6a**). One interface is formed by Oxy2 residues R83 and H84 that are connected to β1 (based on the previously-described secondary-structure^59^) interact with a CaM loop consisting of residues D132 and D134, which coordinate a Ca^2+^ ion (EF4). Thus, R83 and H84 sidechains likely make backbone contacts with this loop. A second interaction region is formed by negatively charged residues (D119, E120, and D123) of CaM H7 and positively charged residues R86, K88 in the Oxy2 β1 and K497 in the Oxy1 H18 (**Fig. 4a (left) and b, I–II**). While the map resolution was insufficient to pinpoint side-chain interactions, the positioning of these residues indicates several potential salt-bridge surface contacts between CaM and the Oxy1-Oxy2 interface.

The FAD-FMN domains make two notable cis and trans surface contacts with the Oxy dimer that likely help position the FMN cofactor adjacent to the heme pocket of Oxy2 (**Fig. 4b, panels III–IV**). We identify long-range cis contacts between the FAD and Oxy1 involving a well-ordered interaction between Y812 at the tip of the β-hairpin in the FAD and R398 in the Oxy1, which is indicative of a potential cation-π interaction and may be critical for stabilizing the FMN-FAD position across the Oxy dimer surface (**Fig. 4b, panel III**). Additionally, V814 of the FAD contacts the aliphatic chain of the Oxy1 R397, potentially forming a hydrophobic interaction, while the FAD K817 forms a putative salt bridge with Oxy1 E402, further stabilizing this site. FAD interactions extend to the Oxy2 domain as well, with loops containing residues N731, N766 and N963 forming potential hydrophobic and hydrogen-bonding contacts with H7 and H13 of Oxy2, which may further stabilize the interface (**Extended Data Fig. 6b)**. We additionally observe potential electrostatic contacts between the FMN and Oxy2 including charged residues (D573, K574, and R576) of the FMN domain, which are positioned adjacent K155 and E156 in the Oxy2 at the tip of a helix-turn-helix motif (HTH) formed by helices H3 and H4 (**Fig. 4b, panel IV**).

To assess the functional significance of these interactions, we performed NOS activity assays, as described above for the wild type (See Methods) (**Fig. 4c**). We note that for each of these mutants, including double and triple combinations, the stability and folding of the iNOS:CaM complex appeared equivalent to wild type, as assessed by comparable purity after tandem Ni-NTA and ADP-resin purification and by the presence of CaM and heme (**Extended Data Fig. 6c,d**). Disrupting the interaction between Oxy2 and CaM by mutating both R83 and H84 to alanine reduced the activity of the iNOS:CaM complex by ∼25%, indicating this interaction, despite being distal from the cofactor sites, is important for iNOS activity, likely through positioning CaM on the Oxy dimer as indicated by the structure (**Fig. 4c**). Strikingly, neutralizing the charged residues of CaM by alanine substitution (D119A, E120A, and D123A), which appear to interact at the Oxy1-Oxy2 interface (**Fig. 4b**), nearly abolishes iNOS:CaM activity (**Fig. 4c**), indicating these residues are indeed critical for CaM-promoted NOS function. Notably we observe no differences in CaM occupancy of the iNOS complex during purification for this mutant (**Extended Data Fig. 6c**). These results are supported by previous studies on CaM activation of NOS, and reveal an interaction mechanism in which these charged residues contact a complementary site at the Oxy1-Oxy2 interface, likely stabilizing the position of the Red arm over the Oxy dimer for FMN-heme electron transfer^26,28^. Individual charge-reversal mutations of the E120-K497 residue pair, previously untested, significantly reduced iNOS:CaM activity by approximately half (**Fig. 4c**). Moreover, combining the two mutations (E120K and K497E) restored activity, thereby establishing the functional significance of this salt bridge interaction, as predicted by our structural model. Interaction III was tested by a Y812A mutation in the FAD domain, which reduced the activity by nearly half, supporting this site in stabilizing the FAD–Oxy1 interaction (**Fig. 4b,c**). Finally, the HTH motif (interaction IV) was tested by mutating residue K155 in Oxy2, but neither K155D nor K155A substantially affected activity (**Fig. 4b,c**). Conversely, a Glu to Arg charge reversal mutation of the adjacent residue (E156R) did reduce the activity of iNOS:CaM by ∼25%, supporting that this residue may help coordinate the FMN domain. Overall, while our mutational analysis did not identify single sites that completely abolish activity, this was expected given the large protein-protein interaction interface we identify, which likely contributes collectively to organize the iNOS engaged state. Taken together, the structure and corresponding mutational analysis identify four key interdomain and interprotomer interaction sites across the iNOS:CaM complex that support a Red^DS^-Oxy-engaged arrangement for FMN-domain positioning and interprotomer electron transfer.

### Conformational flexibility enables FAD–FMN and FMN–heme cofactor alignment

In our iNOS:CaM structure refined from a single well-resolved class, the Red domain remained at a lower resolution compared to the Oxy dimer, indicating flexibility between these regions (**Extended Data Fig. 3a**). Indeed, comparison of Red-containing classes show differences in Red domain position relative to the Oxy dimer, with 8-9Å shifts along the dimer surface, either lengthwise (comparison of Class 1 and Class 4) or laterally (comparison of Class 1 and Class 3) (**Extended Data Fig. 2d and 7**). To further assess these conformations and determine whether the Red domain is continuously flexible or adopts distinct positions relative to the Oxy dimer we further examined the six 3D classes by performing 3D variability analysis (3DVA) to model the Red conformational landscape (**Extended Data Fig. 2d and Supplementary Video 3)**. By 3DVA we identify states with two Red-Oxy positions within the engaged state, termed “distal” and “proximal”, which were further refined to ∼3.0 Å resolution and modeled, as above (**Extended Data Fig. 3b,c and 8a,b)**. While the Red domain remained at a similar lower resolution compared to the Oxy dimer, these structures importantly reveal two distinct positions of the Red domain involving changes in r.m.s.d. of up to ∼12 Å for regions of the FAD and FMN subdomains and CaM that are adjacent to the Oxy dimer surface (**Fig. 5a,b and Extended Data Fig. 8c**).

Remarkably, comparison of the distal and proximal states reveals conformational changes that reposition the FMN relative to the FAD and heme cofactor sites (**Fig. 5c and Supplementary Video 4**). In addition to the cofactors, these changes bring FAD and FMN into closer proximity to neighboring aromatic residues implicated in interdomain electron transfer. In particular, Y631 in the Red domain adjacent the FMN and W372 in Oxy are conserved and facilitate electron transfer between the sites^25–27,60^. Notably, the proximal state reveals additional density for the FMN and Y631 just adjacent to the Oxy2 surface, where a small channel provides direct access to the heme cofactor via W372 (**Fig. 5c, right panel, and Extended Data Fig. 8d**)^25–27^. This density is not observed in the same position in the distal state, thus the FMN and Y631 are further separated from the Oxy2 but appear closer to the FAD (**Fig. 5c, left panel**). To further assess these changes, distances were measured at the shortest interatomic sites, revealing that in the distal state, the FMN cofactor is positioned 11.4 Å from the FAD and 13.4 Å and 10.3 Å away from the heme and W372 in the Oxy2, respectively (**Fig. 5c, bottom panels**). Conversely, in the proximal state, the FMN is shifted to be 12.5 Å from the FAD and 9.1 Å and 5.3 Å from the heme and W372, respectively. While these shifts are modest, the close, ∼5 Å distance in the proximal state between the indole ring of W372 and C7 and C8 methyl groups of the FMN presents an optimal geometry for electron transfer to the Oxy domain^25,26,61^. Given the overall flexibility, additional Red-Oxy positions are likely present. However, we postulate that the proximal state likely captures the structural organization required for efficient FMN-heme electron transfer across the protomers, whereas the distal state or other states with more widely separated domains could favor FAD-FMN electron transfer. These comparisons thus indicate that Red-Oxy conformational flexibility may facilitate distinct steps of electron transfer by repositioning FMN relative to FAD and heme while CaM is bound and stabilizing a Red^DS^ state.

## Discussion

NOS is a homodimer that produces nitric oxide, an important free radical signaling molecule, by a reaction that requires electron transfer from NADPH to FAD and FMN cofactors located in the reductase domain (Red) of one monomer to the heme in the oxygenase domain (Oxy) of the other monomer. Calmodulin (CaM) binding is a prerequisite for NOS activity. We and others have long sought a high-resolution structure of the NOS:CaM complex to understand the mechanism of the electron transfer steps. Here, we present the structure of an active, electron-transfer–competent iNOS:CaM complex that reveals transient structural states after CaM binding that illuminates how FAD, FMN, and heme cofactors dynamically align during electron transfer (**Fig. 6a**). The structure also reveals how CaM-bound iNOS organizes interdomain architecture through previously unrecognized interfaces. Flexibility within the reductase domain allows FMN to adopt multiple positions relative to FAD and heme while minimizing solvent exposure. Overall, our findings provide a structural basis for NO generation and establish general principles for electron transfer from the reductase domain to heme in NOS enzymes. In that the Red domain of NOS has high homology with NADPH-cytochrome P450 reductase, our findings have broad implications for electron transfer to the hemeproteins of the cytochrome P450 superfamily.

**Fig. 6.**
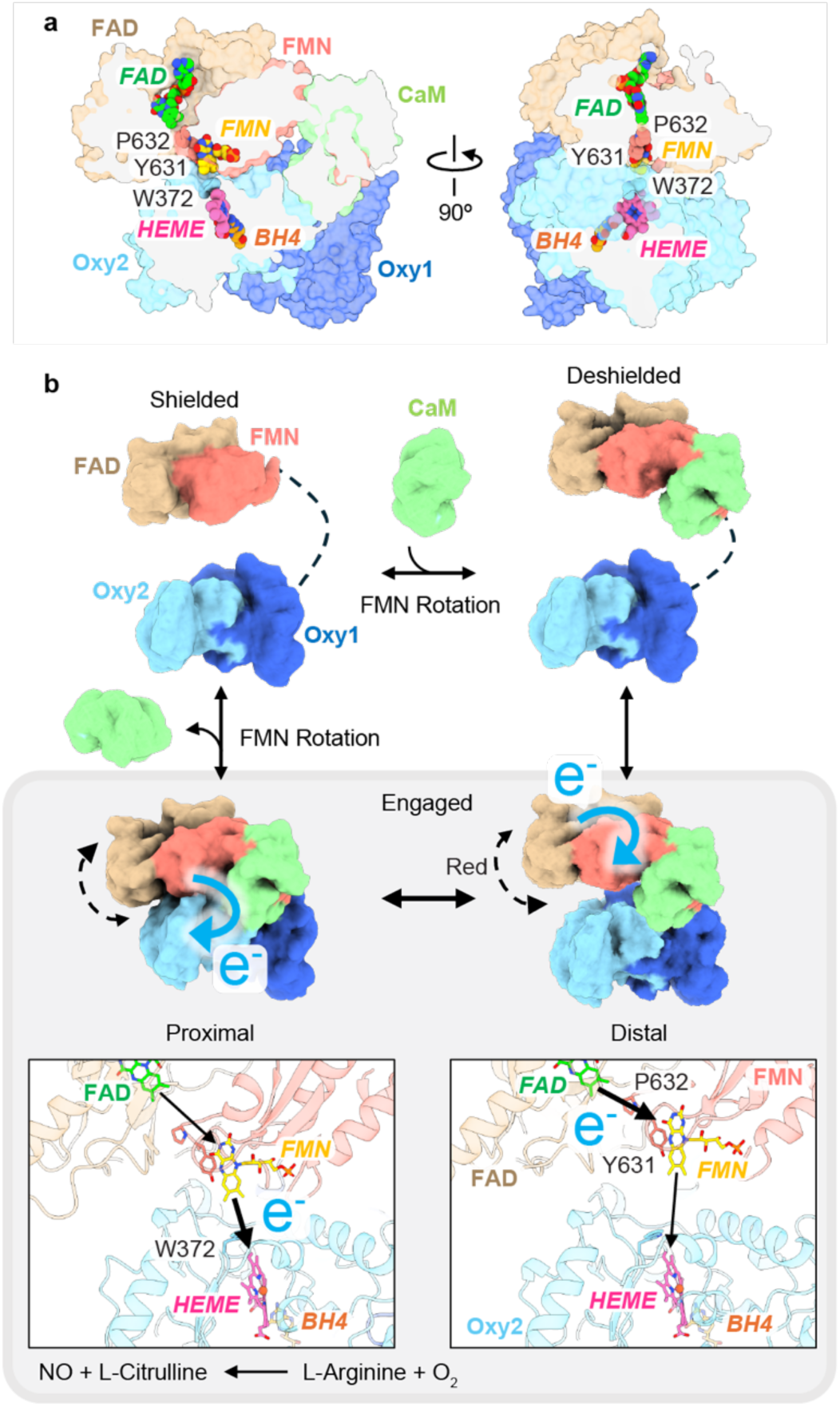
Proposed mechanism and conformational cycle for NO generation. a, Cutaway views of the iNOS:CaM in the engaged state, showing interdomain and cross-monomer alignment of cofactors and conserved residues (shown as spheres) that define a continuous electron-transfer pathway. **b**, Proposed CaM-driven conformational cycle for NO synthesis. CaM binding drives FMN rotation into the deshielded output state, which favors transient engagement of the Oxy dimer surface by the Red domain (gray box) in a conformationally flexible distal state (right) that is more separated from the Oxy heme and may favor FAD–FMN electron transfer. In the proximal state, FMN–heme alignment supports electron transfer to the Oxy domain and NO generation. Electron transfers are indicated by arrows and cofactor positions in the enlarged views of the electron-transfer center (lower panels).

The shielded and deshielded states of the FMN domain have been well-described^36–38^, and it is widely thought that the shielded FMN state allows reduction of the FMN from FAD/NADPH but is protected from redox reactions with other electron acceptors, while the deshielded FMN state moves the FMN away from FAD and closer to the heme to allow for reduction of the heme. The exact structure of the deshielded state has been elusive but is expected to be a state in which FMN is positioned for efficient electron transfer to the heme without uncoupling or electron loss to solvent^26,38,62^. In the current study, we identify a conformational state (proximal state) with an interdomain and interprotomer tunnel in which the FAD, FMN, heme and BH_4_ cofactors are aligned along with key residues (including W372, and Y631 and P632) for efficient electron transfer across ∼5 Å distances (**Fig. 6a**). In this state, the Red domain’s interaction with Oxy protects the FMN from solvent despite the large rotation away from FAD. From our structures we propose a functional model in which iNOS undergoes CaM-mediated, stepwise conformational changes that enable multiple cycles of electron transfer without dissociation of Red-Oxy interface (**Fig. 6b**). CaM binding to the CaM-binding motif induces the FMN de-shielding from FAD. This promotes CaM interaction with the Oxy dimer and FAD engagement with the Oxy domain on the same monomer (Oxy1), positioning the FMN atop the Oxy domain of the neighboring monomer (Oxy2) to form the engaged state. Interestingly, the Red remains flexible in the engaged state, enabling the FMN cofactor to shuttle between FAD-aligned (distal) and heme-aligned (proximal) states. Although it is widely believed that NOS must undergo repeated transitions between a fully shielded and deshielded states for electrons to flow from FAD to FMN to heme, the discovery of distal and proximal states for iNOS:CaM suggests that these states may enable multiple electron transfers in a more rapid manner without large conformational changes. In the case of iNOS where CaM is firmly bound such a mechanism seems advantageous^46,47,63^. It remains to be seen if eNOS and nNOS have proximal and distal states as these isoforms are more dynamically regulated by CaM-binding^46,64^.

In our iNOS:CaM structure, CaM binding to the CaM-binding motif induces de-shielding of FMN from FAD, leading to anchoring of the entire Red^DS^, primarily through the Tyr residue (Y812) at the tip of the β-hairpin finger (or CD2A) of the FAD (**Fig. 4a (right) and b, III**). This beta-hairpin finger motif may serve two roles: (1) as a motif that perturbs CaM interaction under low Ca²⁺ conditions, as previously suggested^65^, while conversely promoting de-shielding of FMN under high Ca²⁺ conditions; and (2) as a structural element that enables interaction of FAD with Oxy and thus facilitates formation of the engaged state. Notably, the β-hairpin finger of neuronal and endothelial NOS is longer than that of inducible NOS, potentially explaining their lower CaM-binding affinity and higher calcium dependency^46,47^. Consistent with this, truncation of the hairpin was shown to reduce the Ca^2+^ dependence of CaM-bound activity of eNOS^65^. In addition, nNOS and eNOS contain the CT and AL elements in Red unlike iNOS (**Extended Data Fig. 5a,f**). It will be important to determine how these additional nNOS and eNOS elements are accommodated in the Red^DS^ state and whether they confer functional roles.

The β-hairpin interaction observed in our studies, together with CaM’s interaction with the Oxy dimer and the hydrophobic FAD–FMN junction, positions FMN atop Oxy2, establishing trans-electron transfer across protomers. The FMN position might be further defined by the HTH motif, uniquely present in the Oxy domain of NOS within the P450 family. The reduced activity caused by the E156R mutation of the HTH may indicate that this motif is required to position FMN on Oxy2. This is also supported by the discrete density of the HTH of Oxy2, observed only when it faces the FMN and not in Oxy1 in the engaged state, indicating a direct interaction, consistent with previous predictions and XL-MS results (**Extended Data Fig. 9**)^28,66^. Together, these extensive and coordinated contacts allow FMN to remain flexible while maintaining structural integrity, potentially enabling multiple rounds of electron transfer by shuttling FMN between FAD and heme without full dissociation.

Several residue interactions change to mediate the conformational shift of Red relative to Oxy (**Extended Data Fig. 8e**). First, in the proximal state, a potential salt bridge between the HTH of Oxy2 (K155) and FMN (D573) is broken as FMN slides toward Oxy2, enabling the formation of new salt bridges by nearby residues E156 and K574/R576 (**Extended Data Fig. 8e, I**). Second, the proximal state may be further stabilized by a possible hydrogen bonding between R452 of Oxy2 and a Ca²⁺-binding loop of CaM (**Extended Data Fig. 8e, II**), consistent with previous molecular dynamics prediction^27^. Finally, the loops containing the FMN cofactor also move toward Oxy2, forming a potential hydrogen bond between E546 of FMN and N208 of Oxy2 and a salt bridge between E551 of FMN and K445 of Oxy2, precisely positioning the FMN cofactor near the heme (**Extended Data Fig. 8e, III**). Supporting these interaction interfaces, previous studies have shown that mutation of E546 disrupts interactions between the FMN and Oxy domains^42,67^, and mutation of the highly conserved residue K445 alters the activity of NOS^31^. Together, the structures presented here reveal how residues at the interfaces mediate interdomain interactions for electron transfer^30,31^.

A methionine residue M434 located adjacent to the critical W372 residue adopts different occupancies in the iNOS:CaM complex and iNOS:CaM Oxy-only densities (**Extended Data Fig. 10**). Previous studies have suggested that a methionine adjacent to a tryptophan residue can facilitate electron transfer by modulating the reduction potential or serving as a “bridging” motif over long distances^68,69^. Thus, M434 may regulate electron transfer and heme accessibility in accordance with Red engagement. In addition, reduced substrate arginine density near the heme was observed for the iNOS:CaM complex in the engaged state, in contrast to the previous nNOS:CaM complex (**Extended Data Fig. 11**). This reduced density may result from NO generation, which is smaller than arginine, prior to sample freezing. Oxy-only densities of the iNOS:CaM complex showed similarly reduced arginine intensity, suggesting that the Red domain could dissociate following NO generation.

Together, this work provides insight into oxygenase–reductase domain interactions in P450 proteins beyond NOS, offering broad implications for understanding how P450s catalyze oxygen insertion into diverse substrates. It identifies conserved features that underlie electron transfer across the P450 family^57,61,70,71^, including FMN dissociation from FAD, orientation of the FMN C7 and C8 methyl groups toward the heme, and adjacent residues that mediate electron transfer. Nevertheless, further studies are required to elucidate their distinct interaction mechanisms with redox partners, as the oxygenase domains of P450s differ from those of NOS^72,73^. Collectively, the results presented here reveal the architecture and dynamics of the iNOS:CaM complex, uncovering a previously unrecognized mechanism of inter-protomer electron transfer underlying NO generation.

## Data availability

The electron microscopy maps and corresponding atomic coordinates have been deposited in the Electron Microscopy Data Bank (EMDB) and the Protein Data Bank (PDB), respectively. The accession codes are as follows: iNOS:CaM Engaged (EMD-XXXXX; PDB XXXX), iNOS:CaM Proximal (EMD-XXXXX; PDB XXXX), iNOS:CaM Distal (EMD-XXXXX; PDB XXXX), and iNOS:CaM Oxy (EMD-XXXXX; PDB XXXX).

## Acknowledgments

We thank the UCSF IND Cryo-EM Facility for assistance with data collection. This work was supported by National Institutes of Health grants GM110001 and GM138690 (to D.R.S.), GM153714 (to Y.O.), and training grant T32GM140223 (to C.M.R.).

## Author Contributions

K.L. designed and performed biochemical and cryo-EM experiments, analyzed data, prepared figures, and wrote and edited the manuscript. C.M.R. and M.L. generated mutant constructs, expressed and purified proteins, and performed enzymatic assays. T.H.P. conducted initial biochemical experiments and cryo-EM data collection. E.T. managed microscopes and the data analysis server. Y.O. supervised the project and provided guidance. D.R.S. designed and supervised the project and wrote and edited the manuscript.

