## Supplementary Materials for "Structural Basis of Electron Transfer by the Human Nitric Oxide Synthase Holoenzyme Complex"

**AFFILIATIONS**

This PDF file includes:

Extended Data Figs. 1–11; Table 1;

Methods;

References for Supplementary Materials

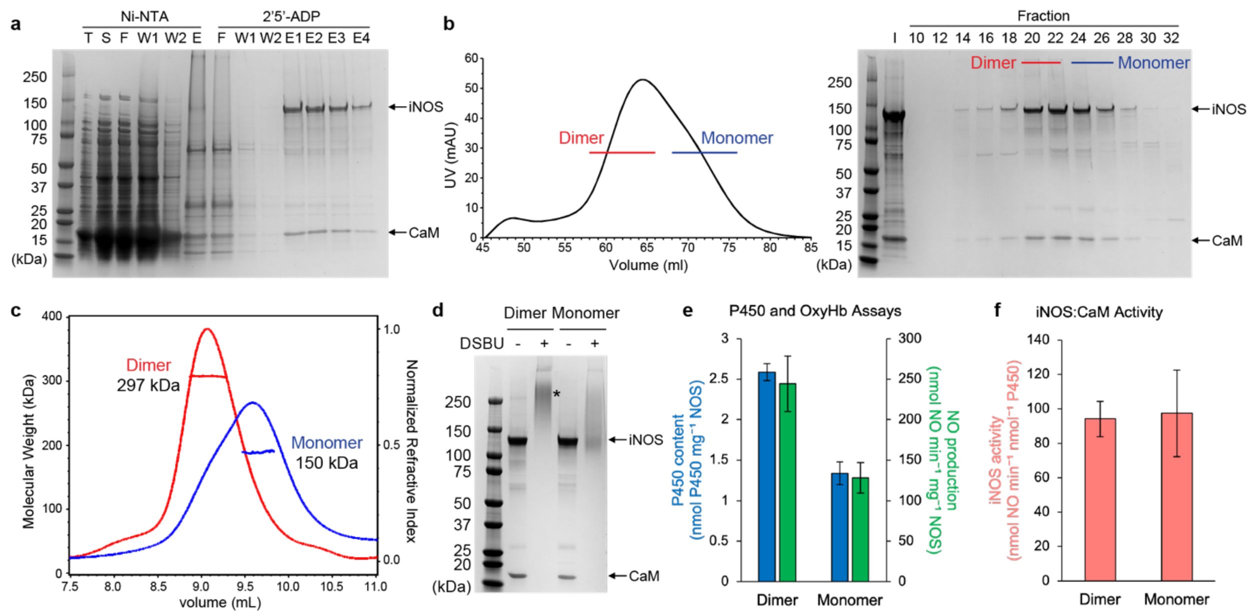


**Extended Data Fig. 1 | Biochemical characterization of the iNOS:CaM complex. a**, SDS-PAGE analysis of iNOS:CaM purified by tandem Ni-NTA and 2',5'-ADP affinity chromatography, with bands corresponding to iNOS and CaM indicated. Lanes: T, total; S, supernatant; W, wash; E, elution; F, flow-through. **b**, Size-exclusion chromatography (SEC) trace of the purified iNOS:CaM complex (left), with SDS-PAGE analysis of the corresponding elution fractions (right). I, Input. **c**, SEC coupled with multi-angle light scattering (MALS) analysis of the purified dimeric (red) and monomeric (blue) fractions of the iNOS:CaM complex, with the corresponding molecular weights determined. **d**, SDS–PAGE analysis of dimeric and monomeric fractions of the iNOS:CaM complex with (+) or without (–) DSBU crosslinking. Although SDS-PAGE detected CaM associated with iNOS in both monomeric and dimeric forms, distinct crosslinked (CXL) species near ~300 kDa were observed predominantly in the dimeric fraction. **e**, P450 concentration (blue), measured by carbon monoxide binding to heme, and NO concentration (green), measured by conversion of oxyhemoglobin to methemoglobin, for the dimeric and monomeric fractions of the iNOS:CaM complex. The residual P450 signal in the monomeric fraction likely reflects incomplete separation of dimers, consistent with the shoulder peak at ~9.0 ml in Extended Data Fig. 1c. As expected, NO production measured by the oxyhemoglobin assay correlated with heme content despite equal protein loading. **f**, Enzymatic activity of dimeric and monomeric iNOS:CaM fractions, measured as NO production normalized to P450 (heme) content, was comparable between the two fractions. Data represent mean ± SD of three technical replicates.

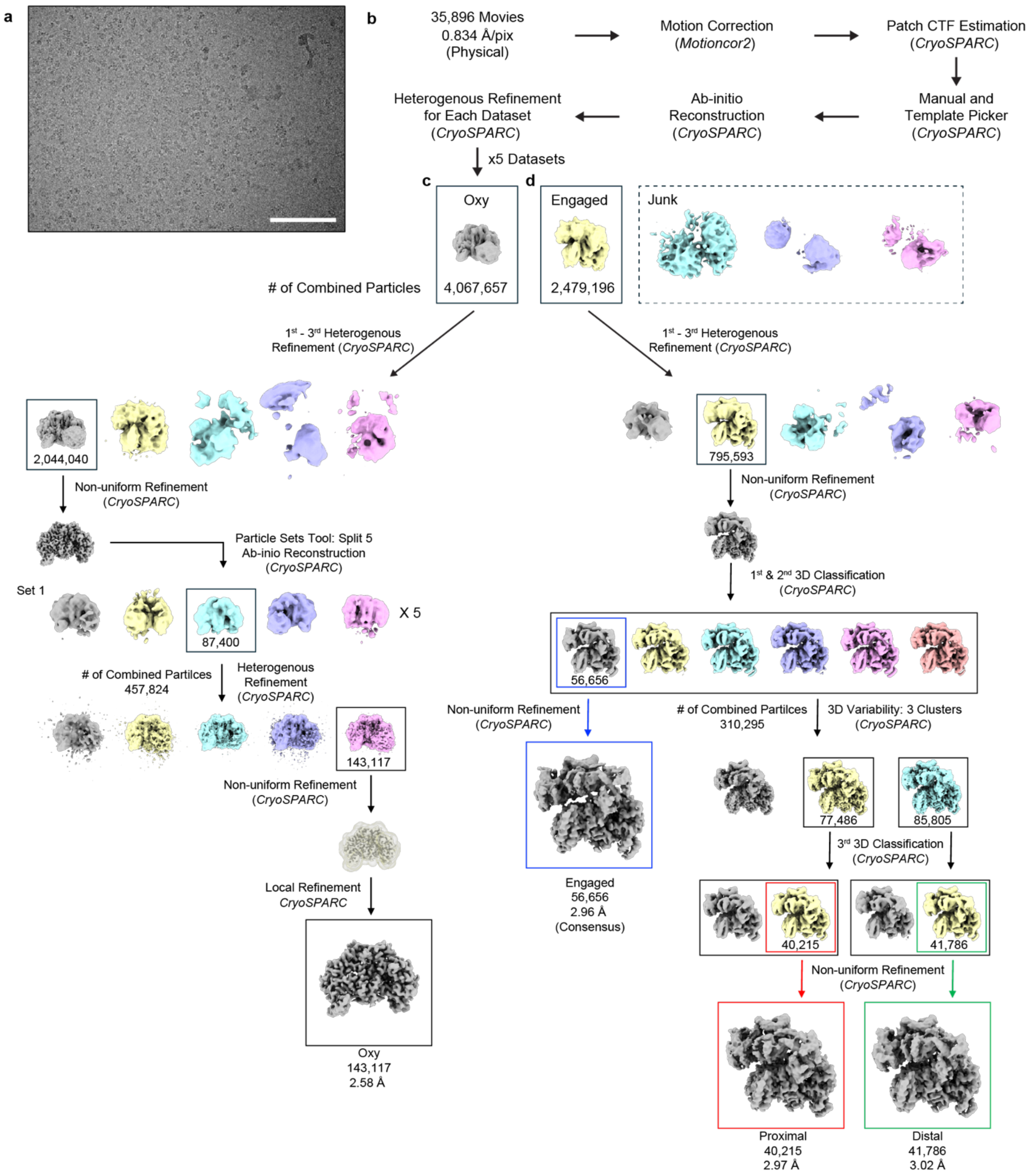


**Extended Data Fig. 2 | Cryo-EM data collection and analysis of the DSBU-crosslinked iNOS:CaM complex. a**, Representative raw cryo-EM micrograph of the DSBU-crosslinked iNOS:CaM complex. Scale bar, 100 nm. **b**, The schematic illustrates the overall workflow for data collection and processing of each dataset (native, DDM, OG, tilted; totaling 35,896 movies)

prior to combination. Standard procedures were followed. Particles were initially picked manually to generate templates for automated picking. The iNOS:CaM engaged and Oxy-only maps were generated via ab initio reconstruction. Three “junk” maps included for heterogeneous refinement throughout the workflow. **c–d**, Data processing for the Oxy dimer with CaM densities (**c**) and the iNOS:CaM complex in the engaged state (**d**), showing the applied processing methods and the classes selected for further refinement. Four final densities were obtained: the Oxy dimer-only density (Oxy) and iNOS:CaM complex densities in the engaged state (Engaged) and its two substates (Proximal and Distal), with corresponding particle numbers and resolutions indicated.

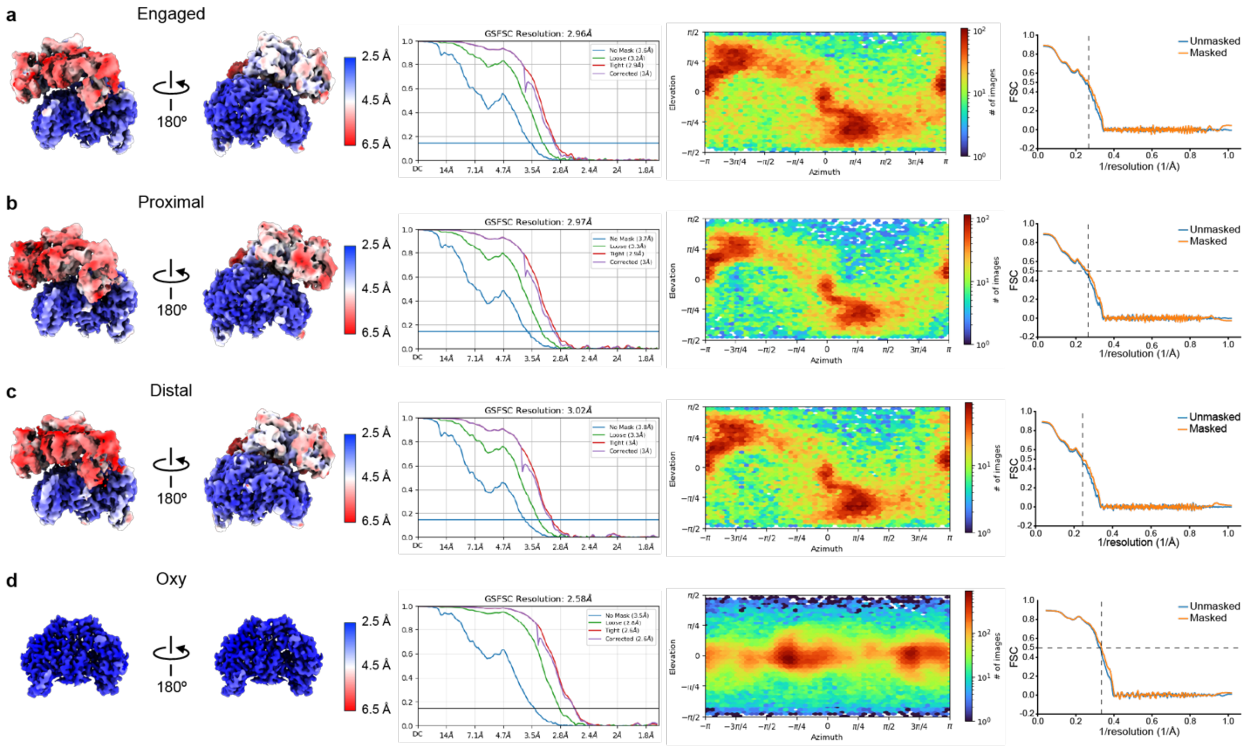


**Extended Data Fig. 3 | Final map and model properties after data processing and molecular modeling.** Final maps with color-coded resolutions, gold-standard Fourier shell correlation (FSC) curves, particle angular distributions, and model-to-map correlations for the cryo-EM structures of the iNOS:CaM complex in the engaged state (**a**), its two substates, proximal (**b**) and distal (**c**), and the Oxy dimer-only density (**d**).

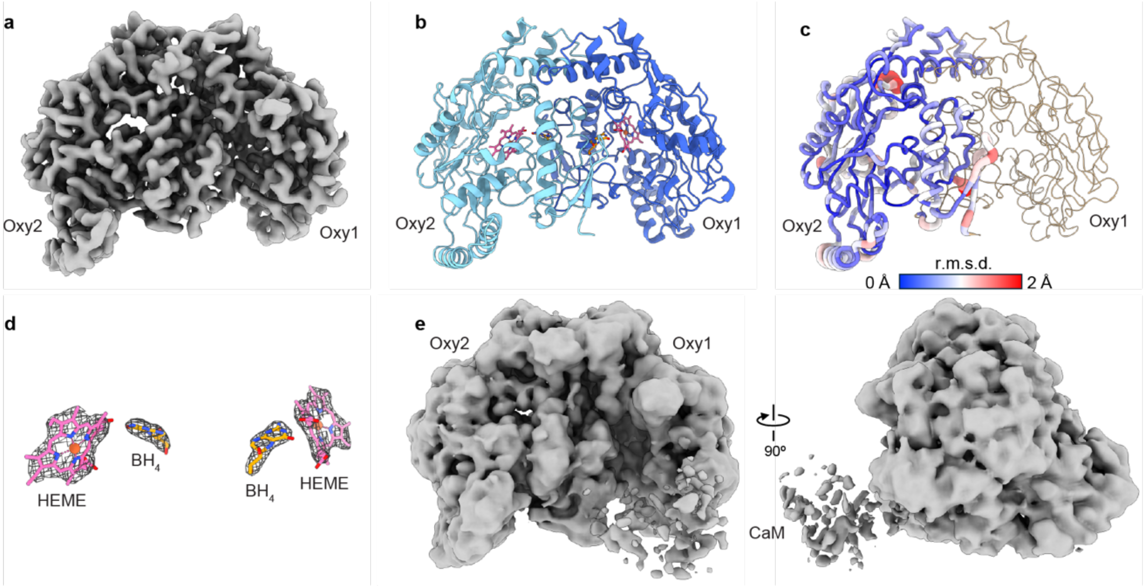


**Extended Data Fig. 4 | Cryo-EM map and model of the iNOS:CaM Oxy-only particles. a**, Refined cryo-EM map of the iNOS:CaM Oxy dimer. **b**, Atomic model corresponding to the refined map. **c**, Cα r.m.s.d. between the iNOS:CaM Oxy cryo-EM structure and the crystal structure of the oxygenase domain of nitric oxide synthase (PDB: 3NQS). **d**, Cryo-EM densities (mesh) corresponding to the heme and $BH_4$ cofactors in the iNOS:CaM Oxy-only complex. **e**, Low-threshold Oxy-only density after Gaussian filtering.

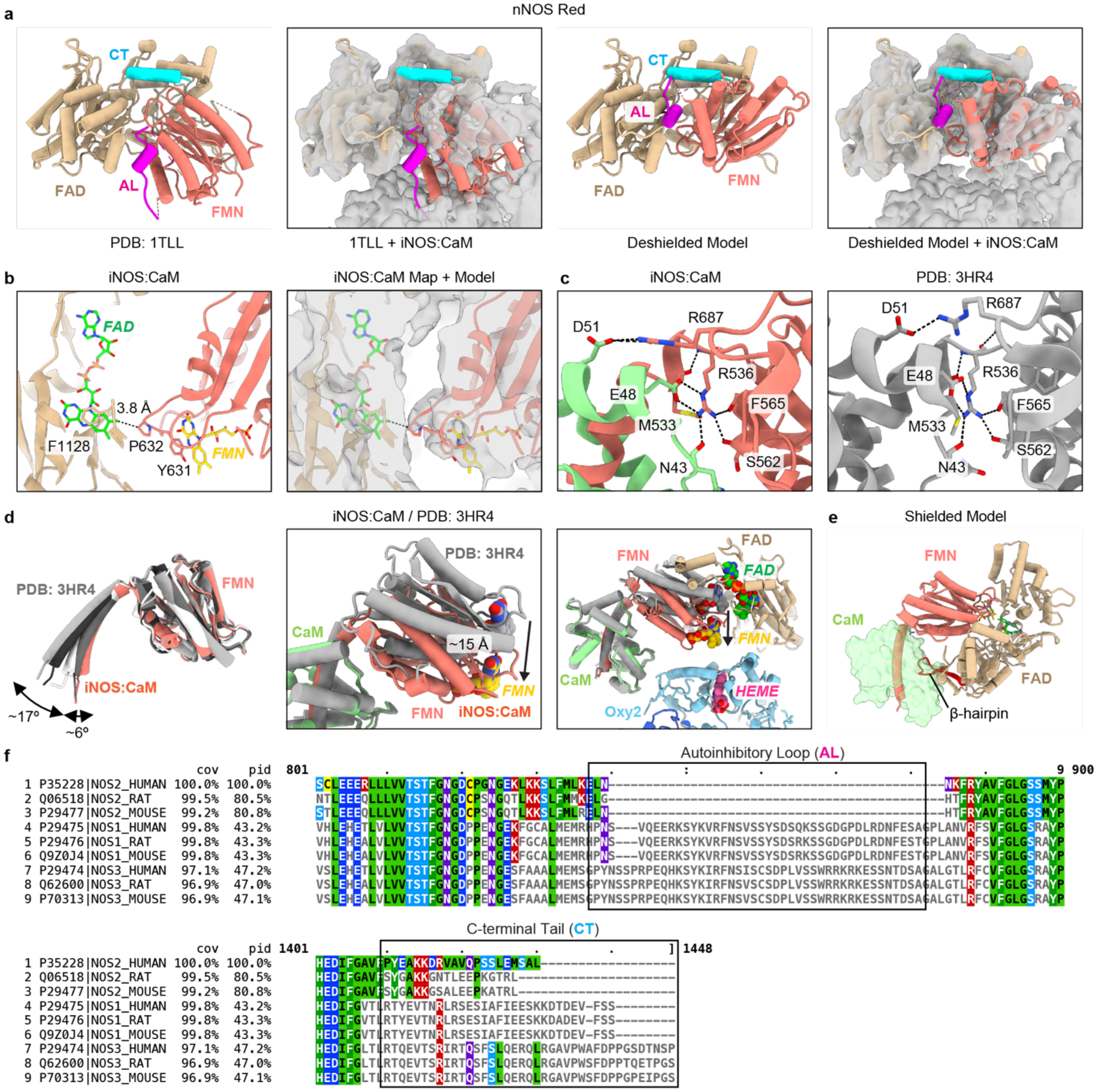


**Extended Data Fig. 5 | FMN subdomain conformations relative to CaM and FAD in cryo-EM and crystal structures. a**. The C-terminal tail (CT) and autoinhibitory loop (AL), which are absent or unresolved in iNOS, are highlighted in cyan and pink, respectively, in the shielded crystal structure (PDB: 1TLL). The deshielded state was modeled based on the cryo-EM structure of the Red conformation of the iNOS:CaM complex. The reductase domain in the shielded and deshielded states of nNOS was docked into the cryo-EM density of the iNOS:CaM complex. **b**, Zoomed-in cutaway views highlight the positions of the FAD and FMN cofactors and adjacent residues shown in stick representation (left), overlaid with transparent gray cryo-EM map (right). **c**, Zoomed-in views highlighting residues coordinating the FMN–CaM interface in the cryo-EM structure (left) and the crystal structure (PDB: 3HR4, right). **d**, Comparison between the cryo-EM and crystal structures of the CaM-bound FMN domain. The FMN crystal structures (PDB: 3HR4, gray) are superimposed onto the cryo-EM structure (colored) (left). A zoomed-in view highlighting the displacement of the FMN cofactor when the CaM-binding motif is used for alignment (middle). A cutaway view showing the positioning of FMN relative to FAD and heme in the deshielded state (right). **e**, The CaM-bound FMN in the shielded state, highlighting a potential clash between CaM (transparent green surface) and the β-hairpin (red cartoon) of the FAD subdomain. **f**, Protein sequence alignment of NOS isoforms from different species. Sequences, indicated by their UniProt accession numbers, were aligned using Clustal Omega. The regions corresponding to the autoinhibitory loop (AL) and C-terminal tail (CT) are highlighted with black boxes.

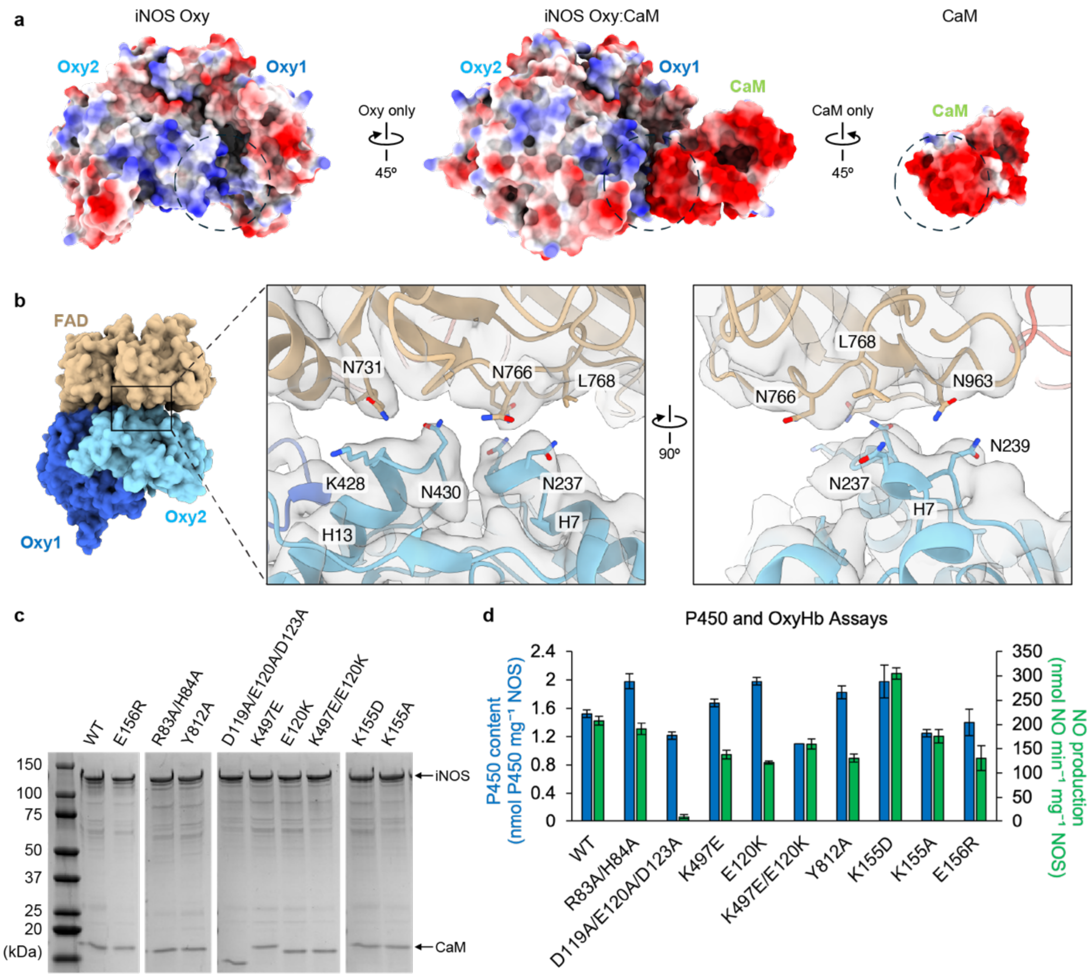


**Extended Data Fig. 6 | Interaction interfaces between the Oxy dimer:CaM and Oxy2:FAD that mediate the engaged state of the iNOS:CaM complex. a**, Electrostatic surface representation of the interface between the oxygenase domain dimer and CaM. Electrostatic potentials of the oxygenase domain of NOS and CaM were calculated and visualized using ChimeraX. **b**, Overall model structure showing the interaction interface mediated by the FAD subdomain and the oxygenase domain 2 (left). Zoomed-in views highlight key residues in stick representation (middle and right), overlaid with transparent gray cryo-EM maps. **c**, SDS-PAGE analysis of wild-type (WT) and mutant NOS:CaM complexes used for P450 and oxyhemoglobin (OxyHb) assays after purification. **d**, P450 (blue), measured via carbon monoxide binding to heme, and NO (green), measured via conversion of oxyhemoglobin to methemoglobin, for iNOS:CaM wild-type (WT) and mutant complexes. Data represent mean ± SD of three technical replicates.

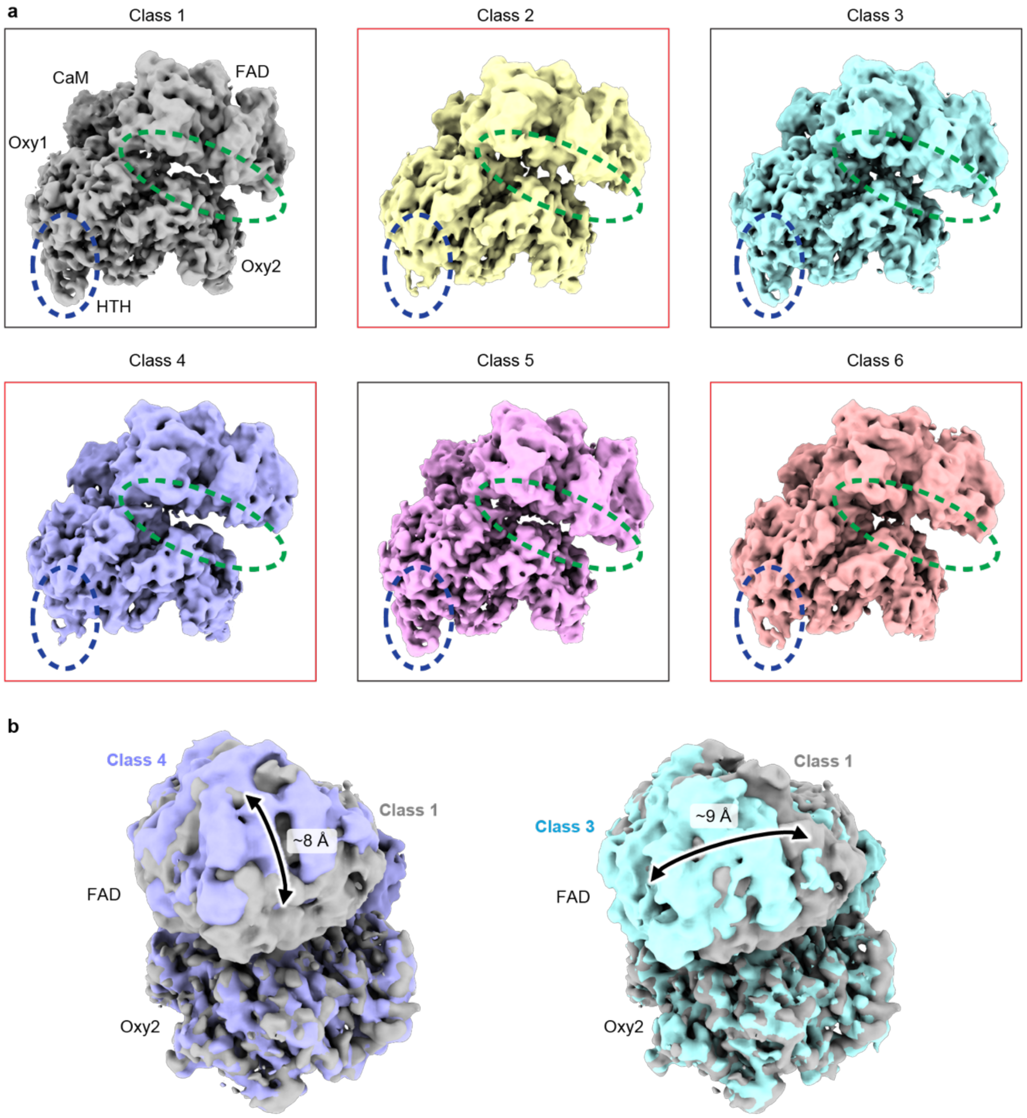


**Extended Data Fig. 7 | Significant conformational flexibility of the Red domain relative to the Oxy dimer in the engaged state of the iNOS:CaM complex. a**, 3D classification of the iNOS:CaM complex in the engaged state from the first non-uniform refinement (Extended Data Fig. 2d) reveals class heterogeneity, primarily reflecting changes in the reductase domain and the helix-turn-helix (HTH) motif of the oxygenase domain. Two distinct subclasses are highlighted with black and red boxes, showing differences in the FAD subdomain position relative to oxygenase domain 2 and the density occupancy of the HTH motif in oxygenase domain 1. **b**, Class 1 (gray) density was globally superimposed with Class 4 (purple, left) and Class 3 (cyan, right) to illustrate vertical and lateral shifts of the FAD subdomain.

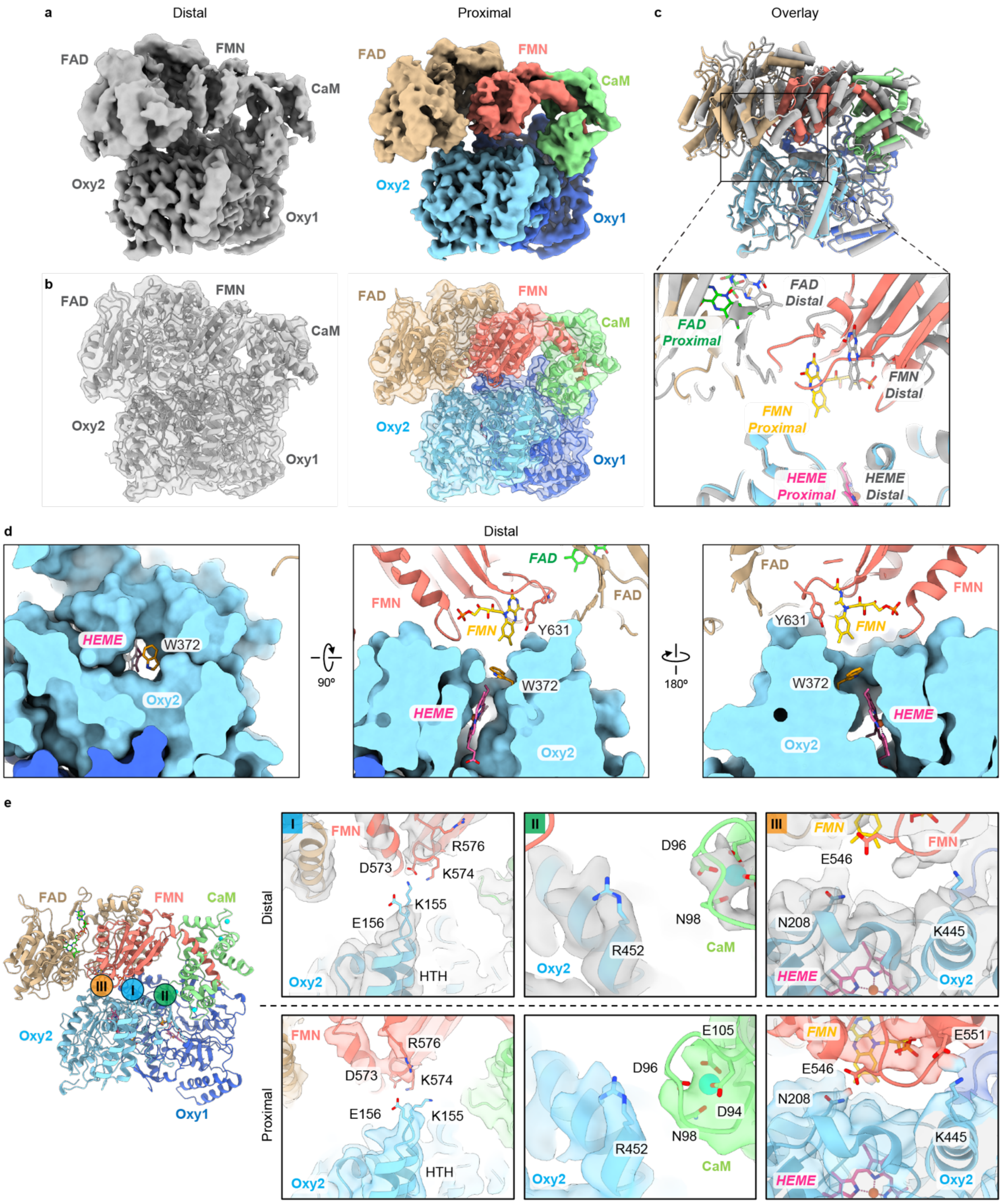


**Extended Data Fig. 8 | The iNOS:CaM complex in distal and proximal states with interface residues. a**, Cryo-EM densities of the iNOS:CaM complex in the engaged state, showing the FMN subdomain relative to the oxygenase domain dimer in distal (gray, left) and proximal (colored, right) positions. **b**, Model structures of the iNOS:CaM complex in distal and proximal states, shown as ribbons, fitted into the corresponding transparent cryo-EM densities. **c**, Superimposed model structures of the iNOS:CaM complex in distal and proximal states, represented as cylinders and stubs in ChimeraX (top), with a zoomed-in view highlighting cofactor positions in stick representation (bottom). **d**, Slab views of Oxy2 in surface representation showing electron transfer tunnel containing W372 and heme (shown as sticks) aligned with the adjacent Red domain FMN cofactor in the proximal state. **e**, Overall ribbon diagram of the iNOS:CaM complex with Roman numerals indicating key interaction interfaces where residue interactions differ between distal and proximal FMN positions (left). Zoomed-in views highlight residues that may interact at the indicated interfaces in stick representation, overlaid with transparent cryo-EM densities in the distal and proximal states (right).

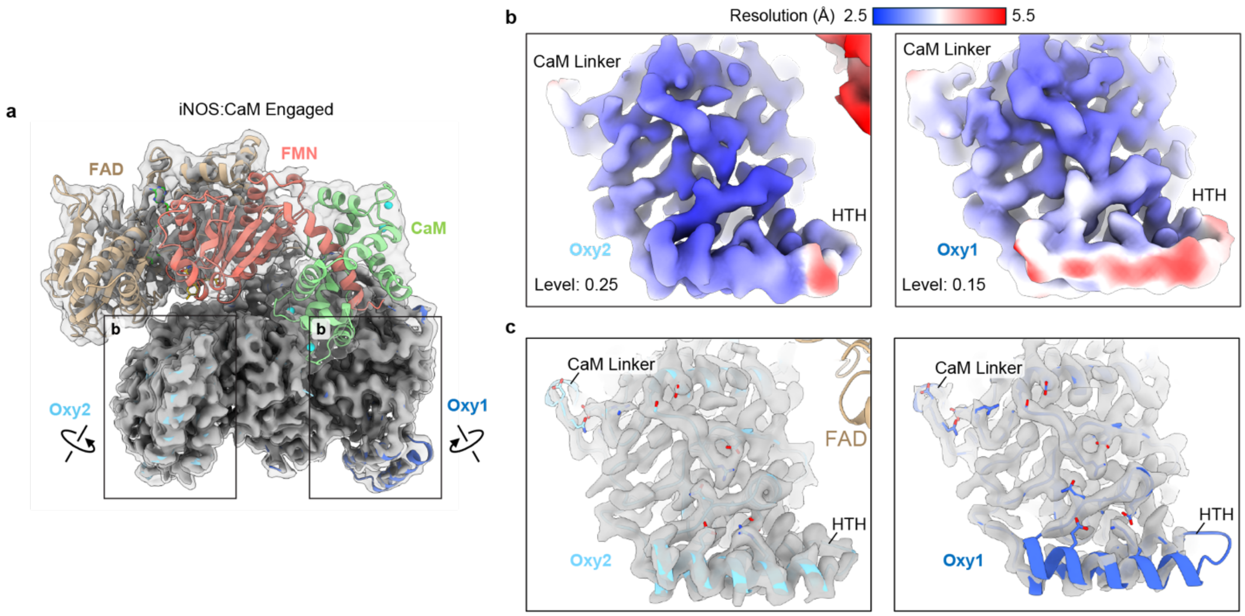


**Extended Data Fig. 9 | The HTH motif interacts more stably with Oxy when oriented toward FMN, with enhanced connectivity extending to the linker connected to the CaM-binding motif. a**, Map and model of the iNOS:CaM complex showing two densities rendered in ChimeraX at thresholds of 0.5 and 0.18 with transparency to highlight differences in the HTH motif (HTH) densities shown in (**b**). **b**, Cryo-EM maps of the HTH motif facing the FMN subdomain (left) and not facing FMN (right), shown at thresholds of 0.25 and 0.15, respectively, to illustrate resolution differences while maintaining comparable density volumes. **c**, Models of the HTH motif and the linker connected to the CaM-binding motif (CaM Linker) in the iNOS:CaM complex facing FMN (left) and not facing FMN (right), overlaid with cryo-EM densities at the same thresholds highlighting key residues shown as sticks that mediate interactions between the CaM Linker and the HTH motif.

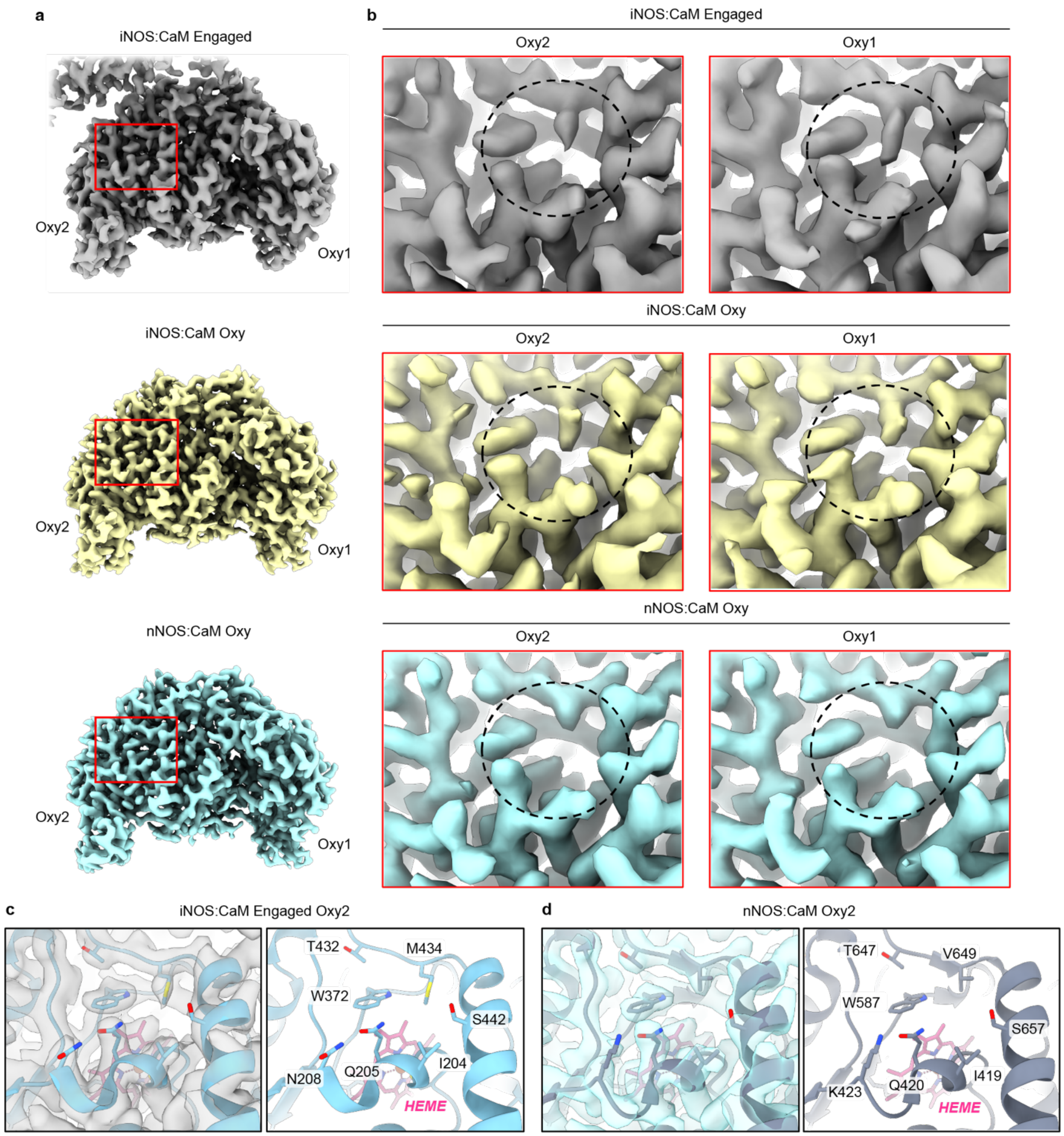


**Extended Data Fig. 10 | Different occupancy of methionine adjacent to the heme, observed in the iNOS:CaM complex in the engaged state, Oxy-only state, and the corresponding residue in the nNOS:CaM Oxy-only state. a**, Cryo-EM densities of the Oxy dimer of the iNOS:CaM complex in the engaged state (top), Oxy-only (middle), and nNOS:CaM Oxy-only (bottom), with a red square indicating the positions of M434 and W372, and the corresponding residues in nNOS, adjacent to the heme. **b**, Zoomed-in views of M434 (V649 in nNOS) densities (black dashed circles) in the iNOS:CaM complex in the engaged state (top), Oxy-only (middle), and nNOS:CaM Oxy-only (bottom). **c–d**, Model structures of M434 and W372 in iNOS and the corresponding residues in nNOS, shown for the iNOS:CaM engaged state (**c**, left) and the nNOS:CaM Oxy-only state (**d**, left), overlaid with transparent cryo-EM densities, alongside the corresponding structures with nearby residue labels (right) adjacent to the heme.

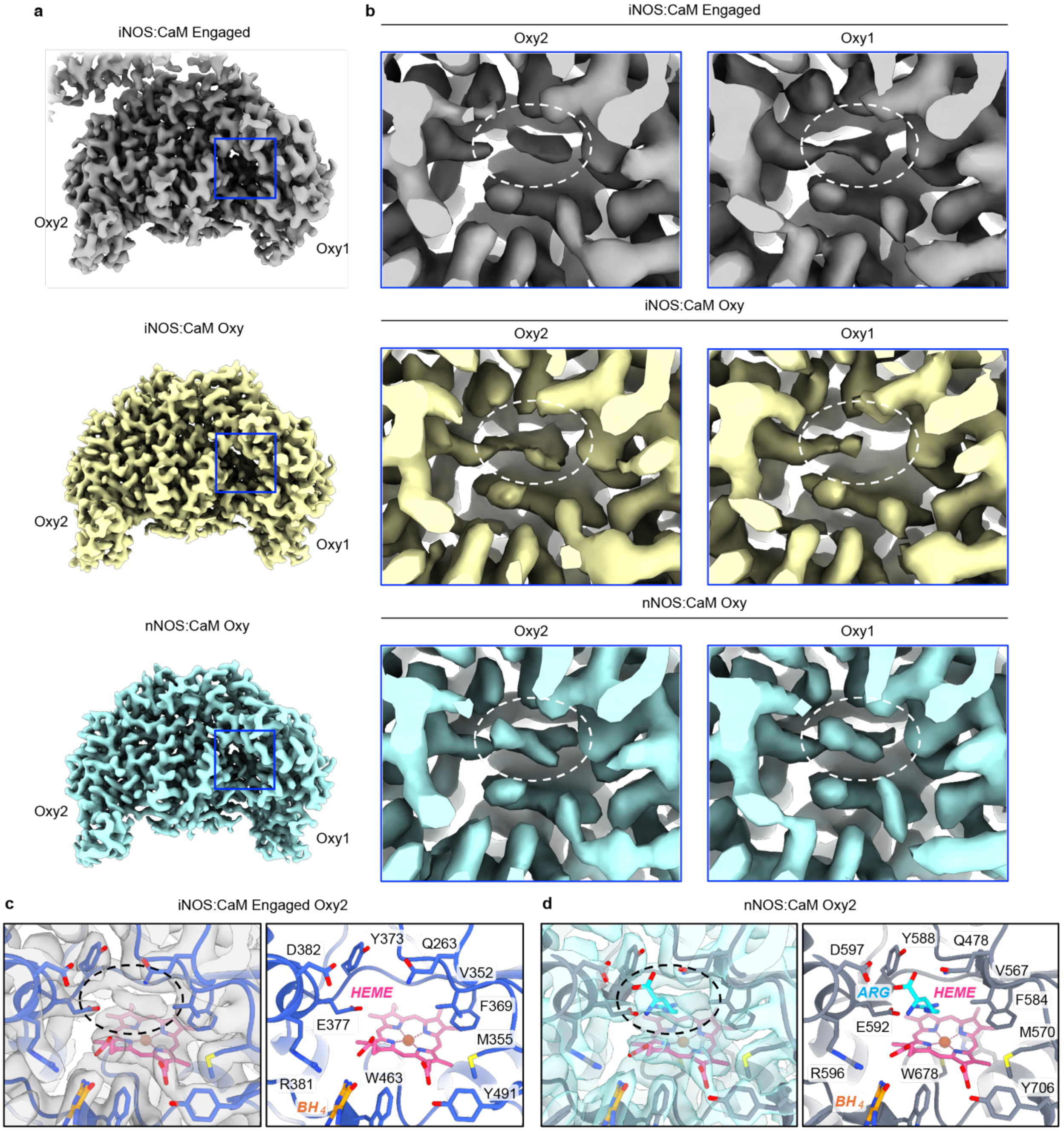


**Extended Data Fig. 11 |Arginine substrate densities observed in the iNOS:CaM complex in the engaged state, Oxy-only state, and in the nNOS:CaM Oxy-only state. a**, Cryo-EM densities of the Oxy dimer of the iNOS:CaM complex in the engaged state (top), Oxy-only (middle), and nNOS:CaM Oxy-only (bottom), with a blue square indicating the substrate-binding site adjacent to the heme. **b**, Zoomed-in views of the substrate-binding sites highlighting substrate densities (white dashed circles) in the iNOS:CaM complex in the engaged state (top),

Oxy-only (middle), and nNOS:CaM Oxy-only (bottom). **c**–**d**, Model structures of the substrate-binding sites in the iNOS:CaM engaged state (**c**, left) and nNOS:CaM Oxy-only state (**d**, left), overlaid with transparent cryo-EM densities, alongside the corresponding structures with residue labels (right). Dashed black circles highlight the substrate densities.

| | iNOS:CaM Engaged | iNOS:CaM Proximal | iNOS:CaM Distal | iNOS:CaM Oxy |
|---|---|---|---|---|
| **Data collection and processing** | | | | |
| Microscope and camera | Titan Krios, K3 | Titan Krios, K3 | Titan Krios, K3 | Titan Krios, K3 |
| Magnification | 59,952 | 59,952 | 59,952 | 59,952 |
| Voltage (kV) | 300 | 300 | 300 | 300 |
| Data acquisition software | SerialEM | SerialEM | SerialEM | SerialEM |
| Exposure navigation | Image shift | Image shift | Image shift | Image shift |
| Electron exposure (e−/Å2) | 66 | 66 | 66 | 66 |
| Defocus range (µm) | −0.8 to −1.8 | −0.8 to −1.8 | −0.8 to −1.8 | −0.8 to −1.8 |
| Pixel size (Å) | 0.834 | 0.834 | 0.834 | 0.834 |
| Symmetry imposed | C1 | C1 | C1 | C1 |
| Initial particle images (no.) | 2,479,196 | 2,479,196 | 2,479,196 | 4,067,657 |
| Final particle images (no.) | 56,656 | 40,215 | 41,786 | 143,117 |
| Map resolution (Å) | 3.0 | 3.0 | 3.0 | 2.6 |
| FSC threshold | 0.143 | 0.143 | 0.143 | 0.143 |
| Map resolution range (Å) | 2.5 – 6.5 | 2.5 – 6.5 | 2.5 – 6.5 | 2.5 – 3.5 |
| **Refinement** | | | | |
| Model resolution (Å) | 3.7 | 3.8 | 4.1 | 3.0 |
| FSC threshold | 0.5 | 0.5 | 0.5 | 0.5 |
| Map sharpening B factor (Å2) | -90.0 | -83.3 | -86.9 | -88.6 |
| Model composition | | | | |

| | | | | |
|---|---|---|---|---|
| Non-hydrogen atoms | 13,207 | 13,207 | 13,207 | 7,085 |
| Protein residues | 1,615 | 1,615 | 1,615 | 854 |
| Ligands | 11 | 11 | 11 | 5 |
| B factors (Å2) | | | | |
| Protein | 79.69 | 79.69 | 79.69 | 82.72 |
| Ligand | 20.74 | 20.74 | 20.74 | 20.08 |
| r.m.s. deviations | | | | |
| Bond lengths (Å) | 0.016 | 0.016 | 0.015 | 0.012 |
| Bond angles (°) | 1.510 | 1.506 | 1.506 | 1.473 |
| **Validation** | | | | |
| MolProbity score | 0.59 | 0.63 | 0.59 | 0.58 |
| Clashscore | 0.23 | 0.35 | 0.23 | 0.22 |
| Poor rotamers (%) | 0.07 | 0.21 | 0.14 | 0.00 |
| Ramachandran plot | | | | |
| Favored (%) | 98.07 | 98.32 | 98.07 | 98.47 |
| Allowed (%) | 1.93 | 1.68 | 1.93 | 1.53 |
| Disallowed (%) | 0.00 | 0.00 | 0.00 | 0.00 |

**Table 1. Cryo-EM data collection, processing, and validation**

### Methods

#### Cloning, expression, and purification of the iNOS:CaM complex

Human inducible nitric oxide synthase (iNOS; NM_000625.4) was cloned into a pTRC-LIC vector (pTRC7; ampicillin-resistant) with an N-terminal 6×His tag using SspI and ligation-independent cloning, omitting the start methionine. Human calmodulin-1 (CaM; NM_001329922.1) was PCR-amplified and cloned into the pCDF7 vector (spectinomycin-resistant) using NdeI and BamHI restriction sites. Site-directed mutagenesis was performed using the Q5 Site-Directed Mutagenesis Kit (New England Biolabs). Protein expression and purification were performed as previously described[1,2] with minor modifications. The two plasmids (pTRC7-iNOS and pCDF7-CaM) were co-transformed into E. coli BL21(AI) cells

(Thermo Fisher Scientific) and selected on LB agar containing the appropriate antibiotics. An overnight culture was used to inoculate terrific broth (12 g tryptone, 24 g yeast extract, 4 mL glycerol, 1 mL of 50 mg/mL spectinomycin, 1 mL of 100 mg/mL ampicillin, and 100 mL of 0.1 M phosphate buffer, pH 7.4, adjusted to 1 L with $ddH_2O$) and grown at 37 °C with shaking at 200 rpm until an $OD_{600}$ of ~0.8 was reached. 5-Aminolevulinic acid hydrochloride (ALA; 1 mL of 0.4 M) was added as a heme precursor, and the culture was incubated at 25 °C with shaking at 120 rpm for 2 h. Expression was induced by adding 1 mL of 1 M IPTG, 2 g of L-arabinose, and 0.5 mL of 0.4 M ALA per flask, followed by overnight incubation at 25 °C with shaking at 120 rpm. Cells were harvested by centrifugation and stored at –80 °C until further use.

Cells were lysed by sonication in lysis buffer (50 mM HEPES pH 7.5, 10% glycerol, 500 mM NaCl, 30 mM imidazole, 1 mM L-arginine, 10 μM tetrahydrobiopterin ($H_4B$), 1 mM phenylmethylsulfonyl fluoride (PMSF), 6 mM β-mercaptoethanol (BME), and Complete Mini EDTA-free protease inhibitor cocktail (Roche)). The clarified lysate, obtained by centrifugation, was incubated with Ni–NTA resin (Thermo Fisher Scientific) for 1 h at 4 °C with gentle rotation, followed by three washes with lysis buffer. Bound proteins were eluted with lysis buffer containing 350 mM imidazole and subsequently incubated with 2′,5′-ADP Sepharose resin (GE Healthcare) pre-equilibrated in ADP buffer (50 mM HEPES, 10% glycerol, 500 mM NaCl, 2 mM $CaCl_2$, 1 mM L-arginine, 10 μM $H_4B$, 0.1 mM PMSF, and 6 mM BME) for 1 h at 4 °C with rotation. After washing with ADP buffer, bound proteins were eluted with the same buffer supplemented with 10 mM $NADP^+$ (RPI).

Eluted proteins were further purified using a Superdex 200 16/600 GL column (Cytiva) on an ÄKTA FPLC system equilibrated with column buffer (50 mM HEPES pH 7.5, 10% glycerol, 100 mM NaCl, 2 mM $CaCl_2$, and 1 mM L-arginine) to separate dimeric and monomeric fractions of the iNOS:CaM complex. Corresponding fractions were pooled, concentrated using centrifugal concentrators, and flash-frozen for future analyses. Proteins used for biochemical assays were concentrated and frozen immediately after $NADP^+$ elution without further purification.

**Biochemical characterization**

Dimeric and monomeric forms of the iNOS:CaM complex were analyzed by size-exclusion chromatography coupled to multi-angle light scattering (SEC–MALS). Complexes (10 μM each) were prepared in running buffer (50 mM HEPES pH 7.5, 100 mM NaCl, 2 mM $CaCl_2$, 10 μM $H_4B$, 6 mM BME, and 1 mM L-arginine) and injected onto a Shodex KW-804 size-exclusion column connected to in-line DAWN HELEOS MALS and Optilab rEX refractive index detectors (Wyatt Technology Corporation) at 4 °C. Molecular weights of the complexes were calculated using the ASTRA V software package (Wyatt Technology Corporation).

For crosslinking, dimeric or monomeric iNOS:CaM (2 μM) were incubated in crosslinking buffer (50 mM HEPES pH 7.5, 100 mM NaCl, 2 mM $CaCl_2$, 10 μM $H_4B$, 10 μM L-arginine, 10 μM NADPH, 10 μM FAD, and 10 μM FMN) for 5 min at room temperature. Disuccinimidyl dibutyric urea (DSBU) was then added to a final concentration of 4 mM, and the reaction was incubated for 20 min at room temperature. Crosslinked samples were quenched with 50 mM Tris–HCl (pH 7.5) and analyzed by SDS–PAGE.

NO synthesis was determined by measuring the NO-catalyzed conversion of oxyhemoglobin to methemoglobin as previously described[3], except 2.0 μg of iNOS was used per assay.

The P450 concentrations were determined by measuring the ferrous carbonyl complex as described previously[4,5] , with the following modifications. The purified WT and mutant iNOS (1 μM) were diluted in a total volume of 100 μL of spectral buffer (0.1 M potassium phosphate, pH 7.4, 20% glycerol, and 0.1 mM EDTA), represented as final concentrations. CO was bubbled through the solution, and a few grains of solid sodium hydrosulfite were added[4]. A spectrum was recorded, and the concentration of the ferrous carbonyl complex of iNOS was calculated using an extinction coefficient of 74 $mM^{-1}cm^{-1}$ at 444 nm, as previously reported[6].

**Cryo-EM sample preparation and data collection**

2 μM of the purified dimeric iNOS:CaM complex was crosslinked with 4 mM DSBU in crosslinking buffer (50 mM HEPES pH 7.5, 100 mM NaCl, 2 mM $CaCl_2$, 10 μM $H_4B$, 10 μM L-arginine, 10 μM NADPH, 10 μM FAD, 10 μM FMN) for 20 min at room temperature following a 5 min pre-incubation. The reaction was quenched with 50 mM Tris-HCl, pH 7.5, for 10 min at RT. After quenching, detergents were added individually (85 μM DDM, 0.01% Octyl β-D-glucopyranoside, 0.05 mg/mL amphipol) as needed to improve angular sampling. Crosslinked samples were applied to either glow-discharged AU 200/300-mesh Quantifoil grids (1.2/1.3) or

graphene oxide grids without glow discharge. Samples were plunge-frozen in liquid ethane using a Vitrobot maintained at 100% humidity and 4 °C after blotting with Whatman filter paper.

Initial screening was performed on a Glacios TEM (Thermo Fisher Scientific) operated at 200 keV, equipped with a Gatan K2 direct electron detector. Data were collected at a calibrated magnification of 53,920x, corresponding to a pixel size of 0.9274 Å/pixel. A defocus range of -0.8–1.8 μm was used, with a total exposure time of 10s fractionated into 100 subframes, yielding a total dose of 62 $e^-/Å^2$ at a dose rate of 5.33 $e^-$/pixel/s. For data collection, a Titan Krios TEM (Thermo Fisher Scientific) operated at 300 keV, equipped with a K3 direct electron detector and a Gatan BioQuantum energy filter (20 eV slit), was used. Super-resolution movies were collected at a magnification of 119,904x (super-resolution pixel size 0.417 Å/pixel) using SerialEM. Defocus values ranged from –2.0 to –1.0 μm, with a dose rate of 8 $e^-$/pixel/s. Each exposure lasted 6 s, recorded in 120 frames at 0.05 s/frame, resulting in a total dose of 66 $e^-/Å^2$. Movies were motion-corrected using MotionCor2[7] and Fourier-cropped by a factor of 2 to the physical pixel size of 0.834 Å/pixel.

**Cryo-EM data processing and structure modeling**

A total of 35,896 movies from native, OG, DDM, amphipol, and GO datasets were collected for the iNOS:CaM complex on a Titan Krios. Each dataset was processed independently in cryoSPARC[8] to isolate the iNOS:CaM complex and Oxy-only densities. CTF estimation was performed with Patch CTF, and micrographs with thick or broken ice, severe contamination, or poor CTF fits were excluded. Approximately 100 particles were manually picked to generate 2D templates for automated particle picking. Reference-free 2D classification and *ab initio* reconstruction revealed two distinct states: the engaged iNOS:CaM complex and the Oxy-only species.

These reconstructions, along with three junk references, were used for heterogeneous refinement without prior 2D classification to minimize selection bias. Particles corresponding to the engaged and Oxy-only states were refined separately. For the engaged complex, iterative heterogeneous refinements and non-uniform refinement were followed by 3D classification into ten classes. Well-resolved classes were reclassified into six classes, and the best subset (engaged, consensus) yielded a 2.96 Å map from 56,656 particles. To probe reductase domain movements, all particles from the second 3D classification were subjected to 3D variability analysis with three clusters, producing 2.97 Å (40,215 particles, proximal) and 3.02 Å (41,786 particles, distal) maps.

For the Oxy-only state, combined particles underwent three rounds of heterogeneous refinement using the initial Oxy-only and engaged-state references with three junk classes. After non-uniform refinement, ~2 million particles were divided into five subsets for *ab initio* reconstruction. The subset displaying discrete Oxy-only density was combined and refined through additional heterogeneous, followed by non-uniform and local refinement using an Oxy mask, yielding a 2.58 Å map from 143,117 particles.

To build a structural model of the iNOS:CaM complex in the engaged state, the Oxy and Red subunit structures predicted by AlphaFold[9] and the crystal structure of CaM bound to FMN (PDB ID: 3HR4)[10] were used as initial models. These structures were docked into the cryo-EM density of the iNOS:CaM complex using the Fit in Map function in ChimeraX[11]. The docked model was subsequently refined using Rosetta FastRelax[12], incorporating cofactors and Zn. The resulting structure was inspected in ChimeraX and further refined in ISOLDE[13]. The final engaged-state model served as a template for building the proximal and distal states, which underwent the same refinement procedures (Rosetta FastRelax and ISOLDE). All models were validated in Phenix[14] and minimally adjusted in ISOLDE and Coot[15] prior to PDB deposition. The Oxy-only density model was built using the Oxy structures predicted by AlphaFold, following the same procedure described above.
